# Bottom-up and top-down control of dispersal across major organismal groups: a coordinated distributed experiment

**DOI:** 10.1101/213256

**Authors:** Emanuel A. Fronhofer, Delphine Legrand, Florian Altermatt, Armelle Ansart, Simon Blanchet, Dries Bonte, Alexis Chaine, Maxime Dahirel, Frederik De Laender, Jonathan De Raedt, Lucie di Gesu, Staffan Jacob, Oliver Kaltz, Estelle Laurent, Chelsea J. Little, Luc Madec, Florent Manzi, Stefano Masier, Felix Pellerin, Frank Pennekamp, Nicolas Schtickzelle, Lieven Therry, Alexandre Vong, Laurane Winandy, Julien Cote

## Abstract

Organisms rarely experience a homogeneous environment. Rather, ecological and evolutionary dynamics unfold in spatially structured and fragmented landscapes, with dispersal as the central process linking these dynamics across spatial scales. Because dispersal is a multi-causal and highly plastic life-history trait, finding general drivers that are of importance across species is challenging but highly relevant for ecological forecasting.

We here tested whether two fundamental ecological forces and main determinants of local population dynamics, top-down and bottom-up control, generally explain dispersal in spatially structured communities. In a coordinated distributed experiment spanning a wide range of actively dispersing organisms, from protozoa to vertebrates, we show that bottom-up control, that is resource limitation, consistently increased dispersal. While top-down control, that is predation risk, was an equally important dispersal driver as bottom-up control, its effect depended on prey and predator space use and whether dispersal occurred on land, in water or in the air: species that routinely use more space than their predators showed increased dispersal in response to predation, specifically in aquatic environments. After establishing these general causes of dispersal, we used a metacommunity model to show that bottom-up and top-down control of dispersal has important consequences for local population fluctuations as well as cascading effects on regional metacommunity dynamics. Context-dependent dispersal reduced local population fluctuations and desynchronized dynamics between communities, two effects that increase population and community stability.

Our study provides unprecedented insights into the generality of the positive resource dependency of dispersal as well as a robust experimental test of current theory predicting that predator-induced dispersal is modulated by prey and predator space use. Our experimental and theoretical work highlights the critical importance of the multi-causal nature of dispersal as well as its cascading effects on regional community dynamics, which are specifically relevant to ecological forecasting.

## Introduction

Dispersal is a life-history trait (Bonte & Dahirel, 2017) that fundamentally impacts spatial population (Hanski & Gaggiotti, 2004) and community dynamics (Vellend, 2010). By linking dynamics between local and regional scales via gene flow, dispersal does not only affect ecology (e.g. Cadotte *et al.*, 2006), but also evolutionary change (e.g., Morgan *et al.*, 2005). Dispersal is especially relevant in the context of current global changes (Urban *et al.*, 2016): Increasingly fragmented landscapes, as well as shifting climatic conditions, may force organisms to disperse in order to survive. However, dispersal is often grossly oversimplified in models Urban *et al.* (2016), a representation which is at odds with the growing awareness that dispersal must be considered in sufficient detail for a better understanding of ecology and evolution as well as for improving biodiversity forecasts Berg *et al.* (2010); Urban *et al.* (2016).

This discrepancy persists because increased realism usually trades off with generality and tractability. Additionally, understanding the causes and consequences of dispersal is challenging, because dispersal is a highly plastic behavioural trait that depends on multiple factors at both the intra- and interspecific level (Clobert *et al.*, 2009; Matthysen, 2012; Legrand *et al.*, 2015). Indeed, empirical approaches have demonstrated that dispersal varies with environmental contexts including resource availability (e.g. Byers, 2000; Imbert & Ronce, 2001; Kennedy & Ward, 2003; Sepulveda & Marczak, 2012; Aguillon & Duckworth, 2015), intraspecific densities (e.g. Matthysen, 2005; Kim *et al.*, 2009; De Meester & Bonte, 2010; Fellous *et al.*, 2012; Kuefler *et al.*, 2013; Bitume *et al.*, 2013; Baines *et al.*, 2014; Pennekamp *et al.*, 2014; Fronhofer *et al.*, 2015b) and interspecific interactions (e.g. Weisser *et al.*, 1999; Hakkarainen *et al.*, 2001; Hauzy *et al.*, 2007; McCauley & Rowe, 2010; Cote *et al.*, 2013; Bestion *et al.*, 2014; Otsuki & Yano, 2014; Baines *et al.*, 2014, 2015; Hammill *et al.*, 2015; Fronhofer *et al.*, 2015a; Tanaka *et al.*, 2016). Theoretical work has investigated causes and consequences of context-dependent dispersal for intraspecific competition (Travis *et al.*, 1999; Metz & Gyllenberg, 2001; Poethke & Hovestadt, 2002), predator-prey interactions (Poethke *et al.*, 2010; Amarasekare, 2007) and species coexistence (Amarasekare, 2010), to name but a few examples.

The challenge is to increase realism by uncovering fundamental drivers of dispersal that are relevant to population and community dynamics while simultaneously avoiding loss of generality and tractability due to a heterogeneous collection of case examples. At the same time, the relative importance of different factors in modifying dispersal across species has yet to be examined. Synthetic datasets which go beyond single species case examples and thereby allow us to resolve these issues and enable us to establish general dispersal patterns are unfortunately lacking (Berg *et al.*, 2010; Urban *et al.*, 2016).

One approach to resolving this problem would be a meta-analysis of published information. However, this approach, based on many studies that each examine a single factor in one specific species, can misestimate the importance of interactions between factors. Few studies manipulate multiple drivers of dispersal at the same time with precludes us from estimating the relative importance of different dispersal drivers and their interactions. Yet, dispersal is likely determined by multiple factors simultaneously and interactions between these factors are predicted to lead to non-linear effects (Bowler & Benton, 2005; Matthysen, 2012; Legrand *et al.*, 2015; Jacob *et al.*, 2016; Cote *et al.*, 2017). An alternative approach to this problem, therefore, is to conduct a coordinated distributed experiment (Lessard *et al.*, 2012; Fraser *et al.*, 2013; Borer *et al.*, 2014, 2017) where the same factors are manipulated across many different species. Such an approach generates new data on factors that drive dispersal, minimizes noise due to differences in methodology which can decrease the ability of meta-analyses to detect across species patterns, and, most importantly, allows the investigation of interaction effects between factors that influence dispersal across species.

As dispersal is generally context-dependent (e.g. Bowler & Benton, 2005; Matthysen, 2012; Legrand *et al.*, 2015) we argue that it is best understood and investigated within the relevant community setting. Consequently, dispersal is likely a function of the fundamental ecological forces that determine local population dynamics. By default, these forces include bottom-up (resource availability) and top-down (predation risk) impacts from species that regulate the focal species demography.

We aim at a general test of both the ubiquity and potential consequences of context-dependent dispersal (CDD) in trophic metacommunities. Firstly, we test existing theory and explicitly highlight the multi-causal nature of dispersal (Matthysen, 2012) by simultaneously focusing on resource availability and predation risk as dispersal modulators. We expect resource limitation to generally increase dispersal rates in order to escape from low fitness environments (Matthysen, 2012). Little consensus exists on the response to predation risk, which has been suggested to vary among species and potentially follow a vertebrate–invertebrate dichotomy (Wooster & Sih, 1995), or to depend on space use behaviour of predators and prey (Poethke *et al.*, 2010). Finally, Hammill *et al.* (2015) suggest that predation risk and intra-specific competition interact to determine the dispersal response.

We assessed these hypotheses and the importance of dispersal modulators in a coordinated distributed experimental approach (Lessard *et al.*, 2012; Fraser *et al.*, 2013; Borer *et al.*, 2014, 2017) involving 7 laboratories across Europe and 21 replicated experimental two-patch metacommunities with focal species ranging from protozoa to vertebrates (Tab. S1). This approach allows us to use a unified experimental design with analogous experimental conditions for all study species, making CDD responses comparable across species, hence allowing for generalizations.

After establishing the ubiquity of bottom-up and top-down control of dispersal, we synthesize our experimental results and advance our general understanding of spatial community dynamics by demonstrating the local and regional consequences of CDD in a simple but general two-patch food chain model. Our theoretical results showcase the relevance of CDD for local and regional dynamics, specifically for the stability (Wang & Loreau, 2014) of metacommunities.

## Materials, methods and model description

### Study organisms

We used 21 focal study species (*Armadillidium vulgare*, *Chilomonas* sp., *Colpidium* sp., *Cornu aspersum*, *Cryptomonas* sp., *Deroceras reticulatum*, *Dexiostoma* sp., *Dikerogammarus villosus*, *Gammarus fossarum*, *Lissotriton helveticus*, *Paramecium caudatum*, *Phoxinus phoxinus*, *Pieris brassicae*, *Pirata latitans*, *Platycnemis pennipes*, *Pteronemobius heydenii*, *Tetrahymena elliotti*, *Tetrahymena pyriformis*, *Tetrahymena thermophila*, *Tetranychus urticae*, *Zootoca vivipara*), including aquatic, terrestrial and aerially dispersing taxa of protists, algae, arthropods, molluscs and vertebrates. Resources and predators of these focal species were chosen based on known natural co-occurrences to allow for the possibility of a common evolutionary history (see Supporting Information for details).

### Experimental setup and treatments

Experiments across all study species followed the same general experimental procedure. We used experimental two-patch systems adapted to each study species (for example, species-specific patch sizes, corridor size and positions) in order for experimental populations to reflect naturally occurring densities and living conditions. Experimental conditions therefore ranged from connected microcosms (Altermatt *et al.*, 2015) to semi-natural connected mesocosms (the Metatron Legrand *et al.*, 2012). All experimental metacommunities were characterized by the presence of a ‘hostile matrix’ connecting the patches, which ensured that inter-patch relocation was indeed dispersal (Bowler & Benton, 2005; Ronce, 2007; Clobert *et al.*, 2012), that is, a change of habitat with potential consequences for gene flow, and not routine foraging movement. See the Supporting Information for details.

We applied a full factorial design crossing two levels of resource availability (RA) and predation risk (PRED). Resources were *ad libitum* (‘standard’ condition; standard RA) or seriously limiting (low RA). Predation risk (PRED) was represented by the presence (yes PRED) or absence of cues (no PRED) belonging to a natural and relevant (i.e., shared evolutionary history) predator of the focal species. Predator cues could be chemical, visual and auditory, depending on the biology of the focal species. We manipulated predator cues instead of the physical presence of predators in order to avoid concurrent effects on population dynamics. The treatments were always applied to one patch (‘origin’) that was initially populated by similar densities of individuals of the focal species for each treatment. The second patch (‘target’) always had reference conditions (standard resources, no predator cues) and was initially empty.

After placing a population of individuals in the ‘origin’ patch, treatments were applied at the beginning of an acclimation phase which took approximately one quarter of the time of the subsequent dispersal phase. During the acclimation phase no dispersal was possible. The absolute time of the acclimation and dispersal phases were adapted depending on the focal species (see Supporting Information). All treatments were replicated 5 times, with the exception of few species where replication was lower or higher due to experimental constraints. For some species, the experimental design included a block, which always included replicates of all treatments and was accounted for in the statistical analysis (see below). The coordinated distributed experiment on the 21 focal species was carried out in 7 different laboratories across Europe (see Tab. S1).

### Data collection

Data on dispersal, that is, the number of residents (individuals in the patch of origin at the end of the experiment) and dispersers (individuals that had left their patch of origin and were in the target patch at the end of the experiment) after the dispersal phase in each replicate, were either collected using video recording and analysis (Pennekamp *et al.*, 2015) or by direct observation. Using data from further analyses or literature surveys (specified in the Supporting Information), we collected species specific information for the focal species, resources and predators including: movement, space use, feeding strategy, body size, predator specialization and focal species escape strategies. The latter information was either used directly or in relevant focal species to predator ratios as potential explanatory variables for understanding the modulators of resource and predator impacts on dispersal (see Tab. S1). All data is available at Dryad (http://dx.doi.org/xxxxx).

### Statistical analysis

All statistical analyses were performed using the R Language and Environment for Statistical Computing (version 3.3.3) and occurred in two steps. We first analysed overall treatment effects on all species together using generalized mixed effects models (GLMM) on numbers of residents versus dispersers (binomial error structure with logit link function; ‘glmer’ function of the ‘lme4’ package using the ‘bobyqa’ optimizer). As random effects we included experimental block within species within taxon. We used taxon as a random effect to account for potential phylogenetic non-independence and included the levels ‘protists’, ‘algae’, ‘arthropods’, ‘molluscs’ and ‘vertebrates’ (see Tab. S1). We further included the laboratory in which the experiment was performed as a random effect in order to account for potential experimenter effects.

Overdispersion was accounted for by additionally including an observation level random effect. Model selection was performed on all models from the full model which included an interaction between resource availability and predation risk to the intercept model using AICc (e.g. Burnham *et al.*, 2011). Besides identifying the most parsimonious model, we also provide information on relative variable importance, which is the sum of AICc weights of models in which the variable of interest occurs.

In a second step, species specific models were used to extract log odds ratios. Subsequently, these log odds ratios were used to determine species specific modulators of the global CDD response. Model structure for obtaining log odds ratios (logORs) of both bottom-up (resource availability) and top-down (predation risk) effects was analogous to the global analysis described above. However, the only potential random effect at the species level was ‘block’. In case the specific experiment did not include a block we used a GLM and potential overdispersion was accounted for by using a ‘quasibinomial’ error structure. We only modelled an additive effect of resource availability and predation risk, as the global analysis suggested the absence of an interaction (see results). We nevertheless provide the analysis of the species level effects based on models including the interaction between the two explanatory variables in the Supporting Information Fig. S2 and Tab.S5–S7. For the subsequent analyses, one protist species (*Chilomonas* sp.) was excluded, as the logOR and the associated errors were meaningless due to zero dispersal in the reference treatment (standard resources, no predation).

The statistical analysis of the species level logORs and potential explanatory variables was executed in a meta-analysis framework in order to account for the uncertainty associated with each species specific logOR (‘rma.mv’ function of the ‘metafor’ package). Again, ‘taxon’ and ‘laboratory’ were included as random effects. Model selection using AICc was performed on the additive models including all possible combinations of explanatory variables, which can be found in Tab. S1). Specifically, we used ‘focal species ID’, ‘relevant taxon’, ‘dispersal mode’, ‘focal species feeding strategy’ and ‘log(focal body size)’ for the effect of resource limitation and ‘focal species ID’, ‘relevant taxon’, ‘dispersal mode’, ‘rel. space use’, ‘predator mobility’, ‘predator feeding strategy’, ‘predator specialization’, ‘escape strategy’, ‘log(focal body size)’ and ‘log body size ratio’ for the effect of predation. For further information see Tab. S1. We included ‘focal species ID’ to test whether the responses were truly species specific, that is, varied idiosyncratically between species, or were more readily explained by other explanatory variables.

We finally analysed whether the logORs representing responses to resource limitation and predation risk were correlated. We used a resampling procedure (10000 samples) to take the error associated with each logOR into account. During each iteration, we fit a linear mixed effects model (LMM; ‘lmer’ function of the ‘lme4’ package) relating a resource limitation logOR to a predation risk logOR each drawn from the Gaussian distribution characterized by the estimate and error (s.d.) of the respective logOR. ‘Taxon’ and ‘laboratory’ were included as random effects. We used the distribution (mean and median) of AICc values resulting from the resampling to select between the intercept, the linear and the quadratic model. The resampling procedure allowed us to obtain a distribution for each model parameter. We then used these distributions to visualize median and 95% percentiles of the model fit.

### A simple two-patch food-chain model with CDD

To illustrate the consequences of context-dependent, more precisely resource- and predation-dependent dispersal, we explored the dynamics of a simple, two-patch food-chain model. The basal resource (*R*) is abiotic and flows in and out of the system at a given rate (*ω*). The focal species (*N*) feeds upon this resource and is itself subject to predation by a top predator (*P*). For simplicity, we assume that both consumers follow a linear, that is type I, functional response (feeding rate *a*) and that only the focal species is able to disperse (dispersal rate *m*_*N*_). The dynamics of this food chain in patch *i* are given by:

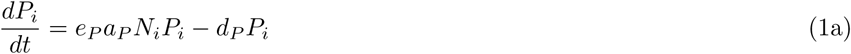

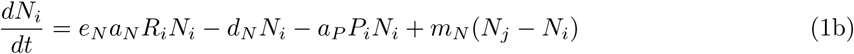

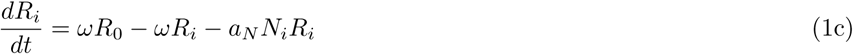

where *e* is the assimilation coefficient, *R*_0_ the resource concentration flowing into the system. The subscripts either indicate the patch (*i*, *j*) or whether the consumer parameters describe the focal species (*N*) or the top predator (*P*).

We compared the dynamics of this two-patch food-chain model with random dispersal (RD) and context-dependent dispersal (CDD). In the earlier scenario, *m*_*N*_ is an unconditional rate. For CDD, we assume that the dispersal reaction norm is a step function as derived by Metz & Gyllenberg (2001). The probability to disperse in the latter scenario will be zero (one) if resources are above (below) a threshold resource density. Simultaneously, the dispersal rate will be zero (one) if predators are below (above) a threshold predator density. In summary, we assume negative resource-dependent dispersal and positive predator-dependent dispersal.

While the RD and CDD scenarios we contrast are characterized by the same model parameters, we compare the specific scenarios in which the RD, respectively CDD, parameters minimize the focal species population dynamics coefficient of variation (CV), as a proxy for local population stability (see also Wang & Loreau, 2014). Alternatively, we compare RD and CDD scenarios that have the same dispersal rates as measured at the end of the analysed time series (see Fig. S4). In analogy to Wang & Loreau (2014), we use temporal coefficients of variation within local communities as well as covariances between communities as proxies for (meta)community stability.

The results we report here should be understood as an illustration of potential consequences of CDD. Although based on a sound mathematical framework (Eqs. 1a–c) and accompanied by a sensitivity analysis (Tabs. S8–S9), the results are a snapshot of possible dynamics as a full analysis of the model is beyond the scope of this manuscript.

## Results

Resource availability and predation risk, that is, the perceived presence of a predator based on chemical, visual and or auditory cues impacted dispersal decisions across all study species (Fig. 1; Tab. S2). The most parsimonious statistical model suggested that the effects of resource availability and predation risk were additive (Tab. S2). While resource limitation led to a clear increase in dispersal across all focal species (on average from approx. 10% to 17% without predation; relative importance of resource availability: 1.00), the effect of predation risk was overall weaker (on average from approx. 10% to 13% without resource limitation; relative importance of predation risk: 0.87). The only marginal improvement of the model including the interaction effect between predation risk and resource availability suggested by the second ranked model (ΔAICc = 1.79; AICc weight = 0.25; see Tab. S2) can be seen when comparing dispersal estimates between the additive and the interactive model in Fig. 1.

**Figure 1:**
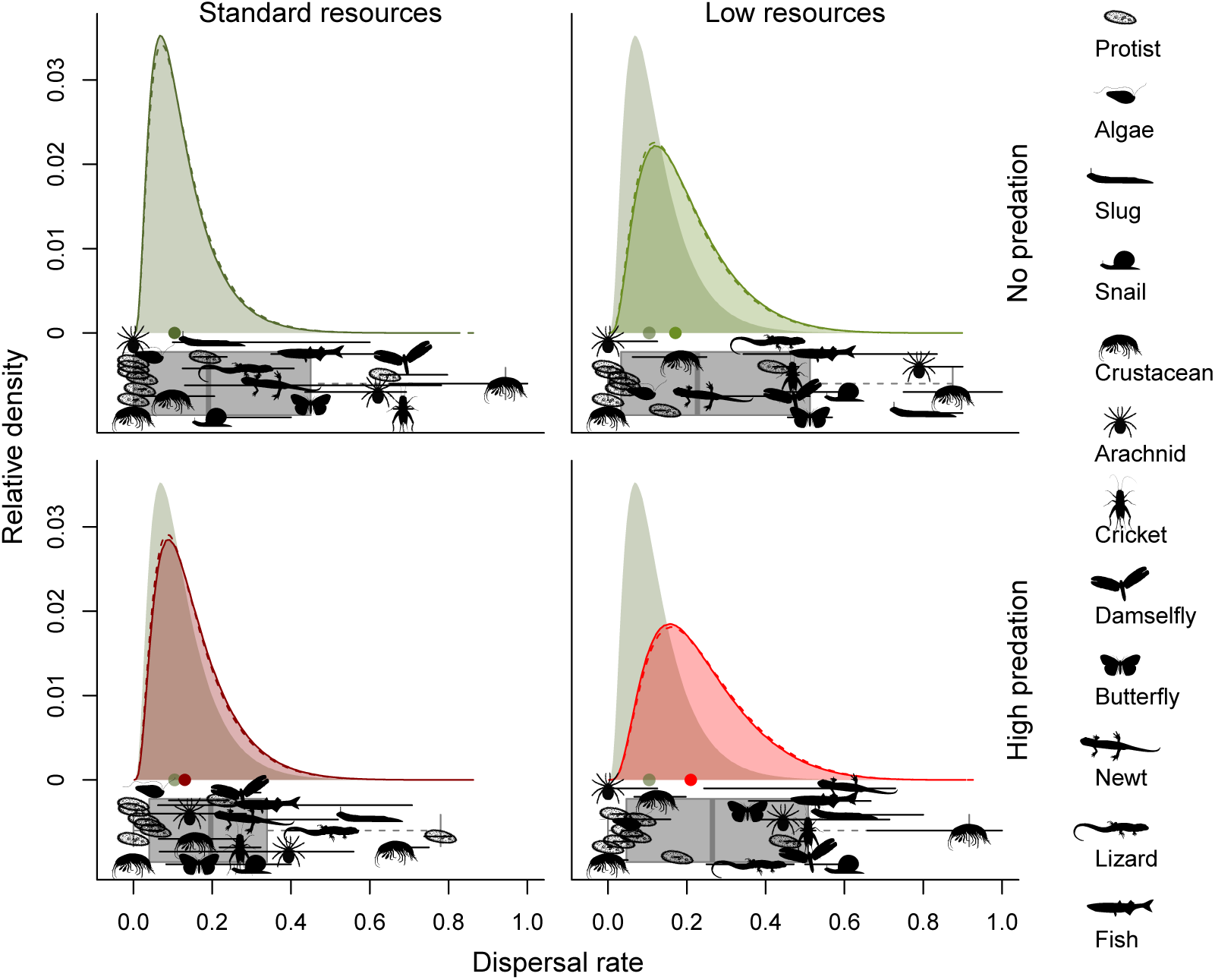
Effect of bottom-up resource limitation and top-down predation risk on dispersal for each study species individually. We here show species trends in dispersal responses (log odds ratios extracted from species specific GLMMs) with respect to species specific dispersal modulators, such as dispersal mode or space use behaviour (see Tab. S1). AICc-based model selection on meta-analytic mixed models confirmed the overall effect of resource limitation (Tab. S3), while the effect of predation risk was modulated by prey and predator properties (Tab. S4). We show dispersal responses (log odds ratios; logOR; black animal symbols) and confidence intervals (vertical black lines) of the resource limitation (top) and predation risk (bottom) effects per species, as well as model estimates (solid coloured lines) and confidence intervals (shaded areas) of the most parsimonious models.

Besides this global analysis, species’ dispersal responses were further analysed using the log odds ratios (logit parameter estimates) extracted from the species specific logistic regression models (Fig. 2). Despite among species variability, the effect of resource limitation was overall positive as in the global analysis: only one species exhibited a clearly negative response to resource limitation (Fig. 2). The response to resource limitation did not depend on species attributes, such as dispersal mode or body size (Tab. S3).

**Figure 2:**
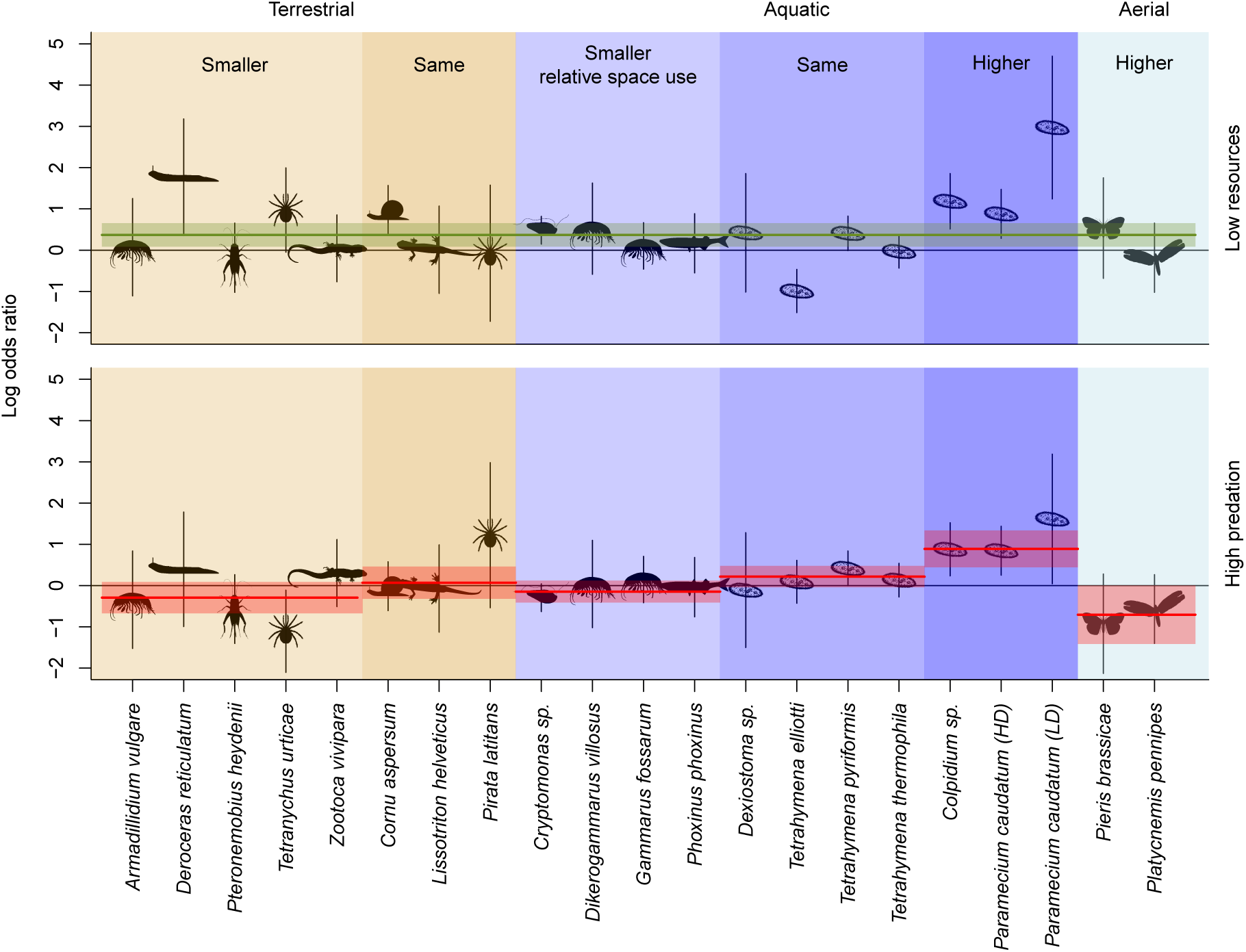
Consequences of CDD for local population and regional metacommunity dynamics. We show the dynamics of all three trophic levels (resources, *R*; focal species, *N*; top predator *P*) in both patches (patch 1: solid lines, patch 2: dashed lines). While the random dispersal (RD) and context-dependent dispersal (CDD) scenarios are characterized by the same model parameters, we compare the specific scenarios in which the RD, respectively CDD, parameters minimize the focal species population dynamics CV, that is, the most locally stable communities sensu Wang & Loreau Wang & Loreau (2014). The RD dispersal rate that minimized the focal species CV was *m*_*N*_ = 0.35. The corresponding CDD thresholds were *T_R_* = 956.94 and *T*_*P*_ = 0.12. Parameter values: ω = 0.5, *R*_0_ = 1000, *e*_*N*_ = 0.1, *a*_*N*_ = 0.01, *d*_*N*_ = 0.1, *e*_*P*_ = 0.005, *a*_*P*_ = 4,*d*_*P*_ = 0.1.

By contrast, the effect of predation risk varied more among species with both positive and negative dispersal responses (Fig. 2). The direction of effects depended on the dispersal mode of the focal species and their relative space use, that is, the extent of space routinely used by the focal species (e.g., a home range) relative to the predator’s space use (Fig. 2, Tab. S4; relative importance of dispersal mode: 0.42; relative importance of space use: 0.33). However, these effects have to be interpreted cautiously, as the second ranking model was the intercept model (ΔAICc = 0.27; AICc weight = 0.17). Further, slightly less important explanatory variables modulating dispersal responses (Tab. S4) were predator specialization (relative importance: 0.36), focal species body size (relative importance: 0.17) and focal species escape strategy (relative importance: 0.16), that is, whether besides dispersal individuals are known to exhibit anti-predator defence, such as hiding or morphological changes. Importantly, as the focal species’ identity was not retained in any of the top models, the responses we report here are not species specific in the sense that they do not vary idiosyncratically between species.

Overall, resource limitation and predation risk were equally important dispersal modulators across all study species: 10 out of 21 species showed a stronger dispersal response to low resources, while 11 out of 21 species showed a stronger response to predation risk. Dispersal responses of species to resource availability and predation risk tended to correlate quadratically (Fig. S3; ΔAICc(linear) = 3.94, ΔAICc(null) = 1.48), implying that species responding to resource limitation by increased dispersal were also likely to respond to predation risk, either by increased or decreased dispersal.

Finally, the two-patch food-chain model shows that the focal species’ CDD may have important consequences for both local as well as regional metacommunity stability (Fig. 3; for a sensitivity analysis see Tabs. S8 – S9). Simultaneous resource- and predator-dependent dispersal reduces fluctuations in local population sizes and thereby increases stability of both focal species and resources strongly (temporal coefficient of variation, CV, of population sizes; see insets of Fig. 3) whereas the effect on the predator’s demography is weaker. Simultaneously, CDD strongly reduces the synchrony of population dynamics between the two patches, thereby also increasing stability (covariation; see insets of Fig. 3). The strong local effects we observe in our model are due to dispersal being simultaneously resource and predator dependent. If CDD is only resource- or predator-dependent, local population fluctuations are reduced to a smaller degree, while the reduction in synchrony may be stronger (see insets of Fig. 3).

**Figure 3:**
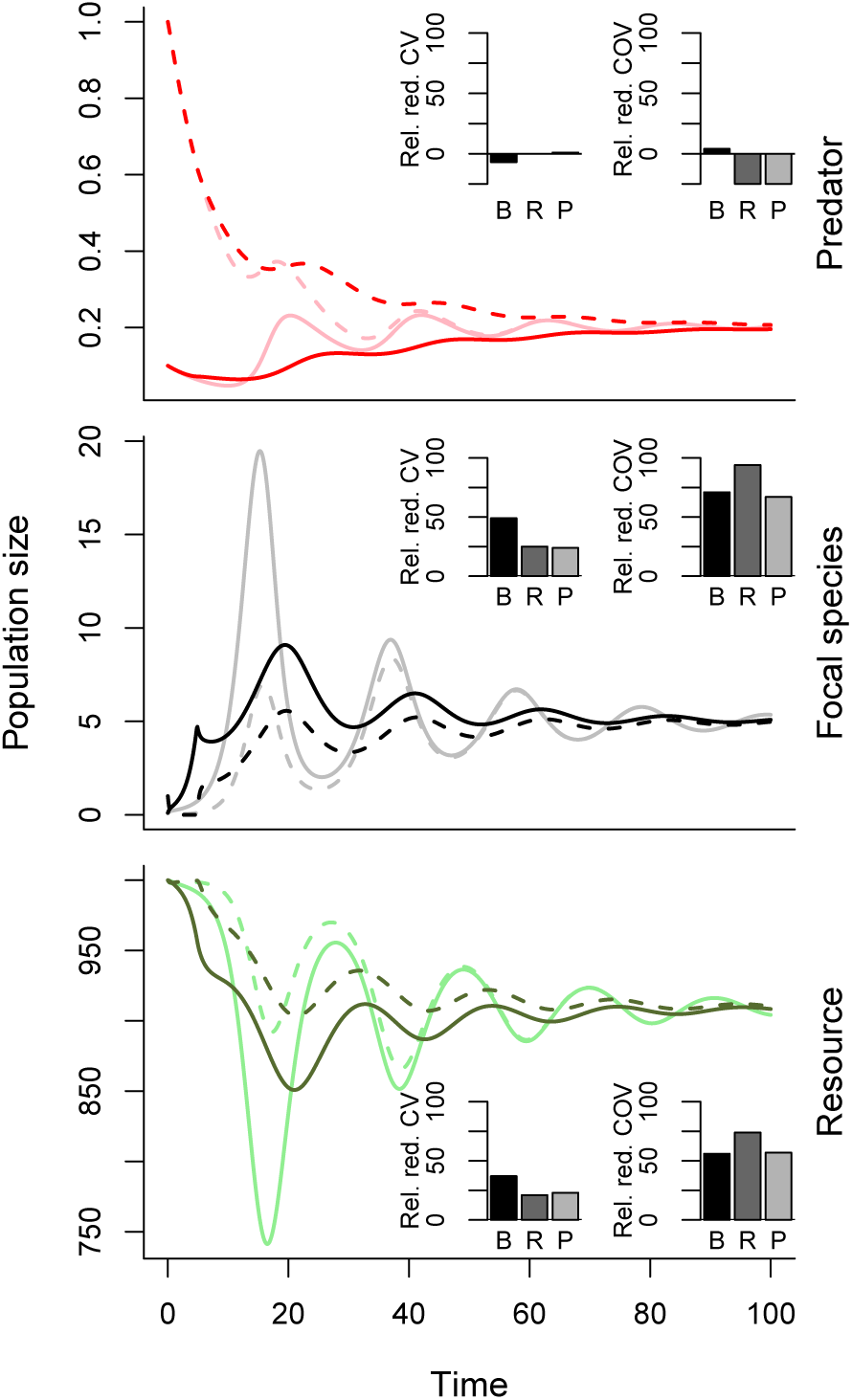
Consequences of CDD for local population and regional metacommunity dynamics. We show the dynamics of all three trophic levels (resources, *R*; focal species, *N*; top predator *P*) in both patches (patch 1: solid lines, patch 2: dashed lines). While the random dispersal (RD) and context-dependent dispersal (CDD) scenarios are characterized by the same model parameters, we compare the specific scenarios in which the RD, respectively CDD, parameters minimize the focal species population dynamics CV, that is, the most locally stable communities sensu Wang & Loreau (2014). The RD dispersal rate that minimized the focal species CV was *m*_*N*_ = 0.35. The corresponding CDD thresholds were *T_R_* = 956.94 and *T*_*P*_ = 0.12. Parameter values: *ω* = 0.5, *R*_0_ = 1000, *e*_*N*_ = 0.1, *a*_*N*_ = 0.01, *d*_*N*_ = 0.1, *e*_*P*_ = 0.005, *a*_*P*_ = 4,*d*_*P*_ = 0.1.

## Discussion

Across 21 focal taxa ranging from protists to vertebrates, we show that dispersal, a central life-history trait (Bonte & Dahirel, 2017) and a regional process, is context-dependent with regards to local bottom-up effects of resource availability, and local top-down effects of predation risk (Fig. 1). While the response to bottom-up effects was a general increase in dispersal (Figs. 1 – 2) top-down modulation (that is, predator-induced dispersal) could be positive or negative (2). Importantly, we show that the latter dispersal responses do not vary idiosyncratically between species but rather depend on the dispersal mode (terrestrial, aquatic or aerial), and on the space use behaviour of prey and predators (Fig. 2).

The differential dispersal responses to bottom-up and top-down forces, which both reduce fitness expectations locally, highlight that local dynamics and species behaviour can only be fully understood if the spatial context is taken into account. In a spatial context, dispersal decisions maximize fitness (Bonte *et al.*, 2014) by balancing (i) fitness expectations in the natal patch with (ii) dispersal costs (Bonte *et al.*, 2012) and with (iii) average fitness expectations in potential target patches. Given that bottom-up and top-down forces both reduce fitness expectations locally, our results suggest that dispersal costs and/or fitness expectations in target patches must differ systematically depending on the type of dispersal trigger (bottom-up versus top-down). This idea has been explored theoretically by Poethke *et al.* (2010), who, very much in line with our empirical results, showed that dispersal responses to predation depend on the spatio-temporal correlation of predation risk, and on the dispersers’ ability to escape from low fitness environments. The modulation of our results by dispersal mode also fits well within this framework, as different dispersal modes imply characteristically different dispersal costs as discussed in Stevens *et al.* (2014). Of course, expanding the range of organisms studied would help determine how general our results are with respect to dispersal mode. In the light of these results, it would be interesting to investigate whether the invertebrate–vertebrate dichotomy in dispersal responses to predation found in streams and discussed by Wooster & Sih (1995) could reflect differences in relative space use.

While both top-down and bottom-up modulators appeared to be equally important across all species (Fig. 2) and the responses tended to covary (Fig. S3), we did not find evidence for a strong interaction between bottom-up and top-down drivers (Fig. 1). This might seem surprising at first, especially given that recent reviews have suggested that dispersal drivers interact (Clobert *et al.*, 2009; Bonte *et al.*, 2012; Matthysen, 2012) and some interactions, especially related to weather conditions, habitat quality and sex, that have been reported empirically (Legrand *et al.*, 2015). However, as dispersal is driven by the above described cost-benefit balance, additive effects are a null expectation while interactions between factors require additional mechanisms that remain to be specified.

Given the general challenges of forecasting ecological dynamics (Petchey *et al.*, 2015; Urban *et al.*, 2016), the absence of a strong interaction has the advantage of making the prediction of ecological metacommunity dynamics potentially easier (Beckage *et al.*, 2011). This finding, along with the general and predictable responses of dispersal to bottom-up and top-down influences, can be seen as encouraging for projecting the dynamics of spatially structured communities into the future. These results could only be achieved using our coordinated distributed experimental approach (Lessard *et al.*, 2012; Fraser *et al.*, 2013; Borer *et al.*, 2014, 2017) using well defined and unified experimental protocols that allow us to achieve generality beyond a meta-analysis which can suffer from little comparability between studies. We here strongly advocate the wide spread use of such large collaborative efforts, as they represent a unique possibility to collect high quality mechanistic data urgently needed for biodiversity forecasting (Urban *et al.*, 2016). Taking the interspecific context, especially multiple trophic levels, and its consequences on dispersal into account will therby be possible and help to better explain and predict spatial community dynamics (Wooster & Sih, 1995).

Shifting our focus from causes of dispersal to its consequences, we demonstrate the important impact of CDD in metacommunities using a simple food chain model (Fig. 3). Our results show that CDD has immediate consequences for local population and regional community dynamics and stability (see also Amarasekare, 2007, 2010; Allhoff *et al.*, 2015). Simultaneous resource and predator-dependent dispersal greatly reduced local fluctuations of population dynamics through time (analogous to *α* variability sensu Wang & Loreau, 2014). At a regional metacommunity level, CDD dramatically reduced covariance between patch dynamics. Both of these effects are directly relevant to metacommunity stability (e.g. Wang & Loreau, 2014), as stability increases with smaller intrinsic fluctuations and less synchronous patch dynamics. These results imply that CDD could mitigate short-term extinction risk in entire metacommunities. Interestingly, CDD in the focal species did not only affect its own dynamics, but had cascading effects on the other trophic levels. Beyond ecological dynamics, plastic dispersal responses have the potential to impact evolutionary as well as eco-evolutionary dynamics as has been recently shown in the context of population differentiation (Schmid & Guillaume, 2017).

In conclusion, our work provides unprecedented insights into the generality of the positive resource dependency of dispersal as well as a robust experimental test of current theory predicting that predator-induced dispersal is modulated by prey and predator space use. Our combination of experimental and theoretical work highlights the far reaching consequences of the multi-causal nature of dispersal as well as its cascading effects on regional metacommunity dynamics, which is specifically relevant to ecological and evolutionary forecasting.

## Author contributions

Author names are sorted alphabetically in the respective sections. All authors commented on drafts and have read and approved the final manuscript.

### Designed the research

Florian Altermatt, Dries Bonte, Alexis Chaine, Julien Cote, Maxime Dahirel, Frederik De Laender, Emanuel A. Fronhofer, Delphine Legrand, Staffan Jacob, Estelle Laurent, Stefano Masier, Frank Pennekamp, Nicolas Schtickzelle. This research was designed during a meeting of the dispNet group (https://dispnet.github.io/) organized at UCL by Nicolas Schtickzelle and Dries Bonte.

### Performed the experiments

Florian Altermatt, Armelle Ansart, Simon Blanchet, Julien Cote, Dries Bonte, Maxime Dahirel, Frederik De Laender, Jonathan De Raedt, Lucie di Gesu, Emanuel A. Fronhofer, Delphine Legrand, Staffan Jacob, Oliver Kaltz, Estelle Laurent, Chelsea J. Little, Luc Madec, Florent Manzi, Stefano Masier, Felix Pellerin, Frank Pennekamp, Nicolas Schtickzelle, Lieven Therry, Alexandre Vong, Laurane Winandy. More information can be found in the Supporting Information.

### Analysed the experimental data

Julien Cote, Emanuel A. Fronhofer

### Designed and analysed the model

Emanuel A. Fronhofer

### Drafted the manuscript

Emanuel A. Fronhofer

## Acknowledgements

FA thanks the Swiss National Science Foundation (Grant No. PP00P3 150698). DB and SM thank the FWO (Fonds Wetenschappelijk Onderzoek — grant n. 11T7518N LV). SJ, EL and NS thank UCL and F.R.S.-FNRS; SJ acknowledges a “Move-In Louvain” postdoc position at UCL; NS is Research Associate of F.R.S.-FNRS. DL and MD thank Fyssen Foundation for funding. JDR thanks the FWO (Research Foundation Flanders - grant n. FWO14/ASP/075). FP was financially supported by Swiss National Science Foundation Grant 31003A 159498. JCo was supported by an ANR-12-JSV7-0004-01, by the ERA-Net BiodivERsA, with the national funder ONEMA, part of the 2012-2013 BiodivERsA call for research proposals and by the French Laboratory of Excellence project “TULIP” (ANR-10-LABX-41).

DL and JC thank Audrey Trochet and Olivier Calvez for their valuable input in experiments involving newts, toads and snakes. MD, AA and LM are especially grateful to Christelle Van Gheluwe for running the experiments, and to Maryvonne Charrier for providing *Deroceras reticulatum* slugs. We thank Mridul Thomas for valuable input on statistical analyses.

## Data accessibility

The data will be made available at Dryad (http://dx.doi.org/xxxxx).

## Supporting Information

Emanuel A. Fronhofer, Delphine Legrand, Florian Altermatt, Armelle Ansart, Simon Blanchet, Dries Bonte, Alexis Chaine, Maxime Dahirel, Frederik De Laender, Jonathan De Raedt, Lucie di Gesu, Staffan Jacob, Oliver Kaltz, Estelle Laurent, Chelsea J. Little, Luc Madec, Florent Manzi, Stefano Masier, Felix Pellerin, Frank Pennekamp, Nicolas Schtickzelle, Lieven Therry, Alexandre Vong, Laurane Winandy and Julien Cote:

## Bottom-up and top-down control of dispersal across major organismal groups

### Supplementary Methods

#### Common woodlouse — *Armadillidium vulgare*

**Authors** Delphine Legrand, Alexandre Vong and Julien Cote

**License** 2012-10 DREAL (common lizard); 09-2016-02 (common toad)

##### Study organism and predator

The common woodlouse (*Armadillidium vulgare*, Latreille 1804) is a widespread European woodlouse species (average body length: 18 mm). In this experiment, we used 236 woodlouse captured in the Metatron (Legrand *et al.*, 2012) and maintained in small tanks (22 L, 39 × 28 × 28 cm). Tanks contained 10 cm of soil litter, 3 egg boxes used as refuges, two small dishes for water and a regular addition of decaying vegetables (apple, grass, potatoes and carrots, Hegarty & Kight 2014). As we caught woodlouse in semi-natural enclosures, they have never been exposed to predator cues (see below).

The common lizard (*Zootoca vivipara*; adult snout–vent length: males, 40–60 mm; females, 45–75 mm) and the common toad (*Bufo bufo*; body length: males, approx. 69 mm; females, approx. 93 mm; Crnobrnja-Isailović *et al.* 2012) were used as our model predator. Both species are generalist feeders preying upon various arthropods species including woodlouse (Avery, 1966; Crnobrnja-Isailović *et al.*, 2012). Lizards were raised in cattle tanks as described in the common lizard protocol and toads were captured in the Metatron.

We estimated that woodlice have smaller home ranges than lizards and toads (< 0.3 m^2^ for woodlice estimated from another isopod species, Hoffmann 1985; < 1200 m^2^ for common lizards, Clobert *et al.* 1994; 8000–28 000 m^2^ for common toads, Parker & Gittins 1979). Mobility, estimated as sprint speed, was higher for lizards and toads than for woodlice (lizard: 30–60 cm s^−1^, Van Damme *et al.* 1991, Sorci *et al.* 1995; common toad: approx. 30 cm s^−1^, Beck & Congdon 2000; woodlouse: 10 cm s^−1^, Dailey *et al.* 2009). On top of dispersal, woodlice have several other anti-predator strategies, including aggregation, sheltering, armour, alarm cue, and repellent chemicals (Broly *et al.*, 2013).

##### Experimental setup

We used 8 dispersal systems placed in a greenhouse with controlled temperature (16–25 °C). Each system is made of two 130 L plastic containers (78 × 56 × 43 cm) connected by a circuitous plastic pipe (diameter: 10 cm, total length: 4.4 m) on the upper section of the container. The departure container was filled with soil litter providing access to the corridor while we only added a thin layer of soil in corridors and arrival containers. To go from the departure to the arrival containers, woodlice had to enter this narrow corridor and fall into the arrival container.

##### Treatments

We created 24 populations made of 8–10 woodlice (approx. 9.83 *±* 0.10 SE). We used a 2 × 2 factorial design, crossing resource availability (RA) and predation risk (PRED) with two levels each, resulting into six replicates of each combination of treatments. We ran the eight replicates in 3 blocks, from September 10th to October 13th 2016. The RA and PRED treatments were applied before releasing woodlice for a 24 hours acclimation phase with connections between containers closed. After 24 hours, connections were opened and we monitored dispersal movements as described below.

##### Resource availability

RA includes two treatments: a low and high resources treatment. The resources are manipulated through the addition of decayed vegetables (half of a potato, half of a carrot, half of an apple and a handle of grass) in the high resources treatment and only a very small piece of vegetable and 2 pieces of grass in the low food resources. It allows creating a large difference in food availability between high and low food enclosures.

##### Predator cues

Lizards were housed in individual terraria containing 3 cm of soil, a shelter (a piece of eggs carton), a water dish, and a piece of absorbent paper to collect odours serving as predator cues. In one corner of the terrarium, ultraviolet and incandescent lamps provided light and heat for thermoregulation from 9:00 to 12:00 and from 14:00 to 17:00. Lizards were fed daily with 1 cricket (*Acheta domestica*). We used a mix of several absorbent papers belonging to several lizard terraria. Four toads were maintained all together in a large plastic tank (130 L, 78 × 56 × 43 cm) with rocks, water, soil and absorbent paper. Tanks were watered daily and abundantly to maintain enough humidity. Before releasing woodlouse, dispersal systems with predator cues treatments received a piece of lizard paper mixes and a piece of toad paper, whereas control treatments received a piece of odor free paper collected from vacant containers maintained under the same conditions than inhabited terraria. When removing a paper from predator or predator-free container, we added a new piece of paper for future experimental blocks. The protocol to collect lizard olfactory cues is a standard protocol already used several times to elicit social reactions in lizards (e.g. Cote & Clobert, 2007; Teyssier *et al.*, 2014) and the woodlice are known to react to predator chemical cues (e.g., volatile ant cues, Hegarty & Kight, 2014). We also added two individuals that were crushed in a tube in each predator treatment as alarm cues of predation (Broly *et al.*, 2013) and an empty tube in no predation treatments.

##### Dispersal and data collection

After the opening of corridors, we monitored dispersal daily for 15 days. To do so, we captured woodlice in the arrival containers without disturbing the departure containers. At the end of the dispersal assay, we captured all woodlice in departure and arrival container. We recaptured 114 of the 236 released woodlice. We only used recaptured ones in our analysis to prevent confounding dispersal with recapture probability.

#### Protists — *Chilomonas* sp., *Colpidium* sp., *Tetrahymena pyriformis* and *Desxiostoma* sp

**Authors** Emanuel A. Fronhofer, Frank Pennekamp and Florian Altermatt

##### Study organisms and predators

Protist microcosms have a long tradition of being used in ecological and evolutionary studies (Altermatt *et al.*, 2015). We here used a set of four predator-prey species pairs cultured in protist pellet medium (0.46 g L^−1^; Carolina Biological Supply) with a mix of three bacterial species as a basal resource (*Serratia fonticola*, *Bacillus subtilis* and *Brevibacillus brevis*). The four species pairs included the focal species *Chilomonas* sp. (body size, as length along the major body axis: 29.7 µm; swimming speed, as mean net speed: 4.72 µm s^−1^) with *Blepharisma* sp. (body size: 91.61 µm; swimming speed: 6.94 µm s^−1^) as a more mobile predator, the focal species *Colpidium* sp. (body size: 75.4 µm; swimming speed: 15.59 µm s^−1^) with the sessile predator species *Stentor* sp. (body size: approx. 800 µm), *Tetrahymena pyrformis* (body size: 37.83 µm; swimming speed: 9.22 µm s^−1^) as a focal species with *Cephalodella* sp. (body size: approx. 300 µm; swimming speed: approx. 10 µm s^−1^), a rotifer, as a slightly more mobile species and finally the focal species *Dexiostoma* sp. (body size: 36.31 µm; swimming speed: 13.16 µm s^−1^) with the less mobile predator *Dileptus* sp. (body size: approx. 200 µm; swimming speed: approx. 8 µm s^−1^). All trait and movement information reported here was measured during the experiments. All protist cultures were originally obtained from Carolina Biological Supply. All focal species graze on the bacterial resource and all predators are generalists and exhibit an active capture strategy, except for *Stentor* sp. which is sessile and therefore a sit-and-wait predator.

##### Experimental setup

We used replicated 2-patch metapopulations consisting of two 20 mL vials (Sarstedt) connected by silicone tubing (VWR) as previously used in (Fronhofer *et al.*, 2015) or (Fronhofer *et al.*, in press). While the inside diameter of the connecting tube was fixed to 4 mm, the length was adjusted to control for different dispersal capacities among focal species: *Chilomonas* sp. and *Dexiostoma* sp.: 7 cm; *Colpidium* sp. and *T. pyriformis*: 3.5 cm.

At the beginning of the experiment the tube connecting the two patches was filled with water instead of protist medium as a ‘hostile matrix’ preventing cell growth and hence reproduction and clamps were used to close the connection. After a 1 h acclimation phase, the clamps were removed and dispersal was allowed for 4 h. All predator-prey systems were replicated six times.

##### Treatments

The composition of the patch of origin depended on the treatment combination which is described below (total volume: 15 mL). By contrast, the target patches all received 13 mL of fresh, sterile medium as well as 2 mL bacterial culture at equilibrium (approx. 1 week old culture) as food resources for the focal species.

##### Resource availability

Resource availability (RA) in the patch of origin was either equal to the target patch (high RA), that is, the patch received 2 mL bacteria at equilibrium (high RA), or severely limited by not providing any bacterial food (low RA: 2 mL water instead of bacteria). To these 2 mL we added 3 mL of a concentrated culture of the focal species. In order to obtain this concentrated culture, we used 15 mL of a culture of the focal species at equilibrium, centrifuged it (Sigma 3-16PK centrifuge; 5 min at 4500 rpm) and washed out the pellet containing the protist cells with 3 mL fresh and sterile medium.

##### Predator cues

The remaining 10 mL of the starting patch consisted either of fresh and sterile medium (no PRED) or sterile filtered medium (filter pore size: 0.2 µm) from the respective predator culture at equilibrium (PRED). The sterile filtered medium from the predator culture contained no bacterial resources but all chemical compounds secreted by the predators which may be used as a predator cue by the focal species. This procedure guaranteed that, regardless the concentration of resources or predator cues (see below) the focal population in the starting patch was always at equilibrium, as the total volume was again 15 mL.

##### Dispersal and data collection

After 1 h acclimation phase and 4 h dispersal we closed the clamps and took 20 s videos in start and target patches using a Leica M205 C stereomicroscope at a 16-fold magnification and a Hamamatsu Orca Flash 4 video camera (imaged volume: 34.4 µL; height: 0.5 mm). Population densities of residents and dispersers were obtained using video analysis and the ‘BEMOVI’ package for the R Language and Environment for Statistical Computing (Pennekamp *et al.*, 2015). For specific settings see GitHub (https://github.com/efronhofer/analysisscriptbemovi).

#### Garden snail — *Cornu aspersum*

**Authors** Maxime Dahirel, Armelle Ansart and Luc Madec

##### Study organism and predator

*Cornu aspersum* (Müller) (Gastropoda, family Helicidae) is a medium-sized land snail common throughout Western Europe (Welter-Schultes, 2012). In May 2016, we collected individuals by hand in suburban populations in Pacé, France (48°9^*′*^ N, 1°47^*′*^ W). Although subadults are overall more dispersive (Dahirel *et al.*, 2016), we selected only adults and old subadults (snails with diameter greater 25 mm) for our experiments, to limit the influence of variation in internal developmental stage. Snails were individually marked with felt-tip paint markers and maintained under controlled conditions (20 °C *±* 1 °C; 16L: 8D). They were housed by random groups of 10 individuals in 30 × 30 × 8 cm polyethylene boxes with 1 cm of moist soil at the bottom. This density is well within the range of naturally observed densities near shelters, and below the thresholds at which many negative effects of crowding appear under controlled conditions (Dan, 1978). Snails were fed ad libitum with commercial snail food (cereal flour supplemented with calcium, Hélinove, Le Boupère, France) placed on a Petri dish at the centre of the box.

Larvae of the European common glow-worm, *Lampyris noctiluca* (Linnaeus) are specialist predators of soft bodied invertebrates, especially land snails and slugs (Symondson, 2004). They are one of the few insect species recorded to predate *Cornu aspersum* snails, and seem to be able to follow snail mucus trails to reach their prey (Symondson, 2004). Crawling glow-worm larvae are about twice as fast on average than crawling garden snails (roughly 4–6 cm min^−1^ versus 2–3 cm min^−1^, De Cock 2009, Dahirel *et al.* 2015). We collected glow-worm larvae by hand in April 2016 in Arçais, France (46°17^*′*^13^*″*^ N, 0°40^*′*^3^*″*^ W). Due to difficulties in maintaining them in controlled conditions, they were killed within a week of capture, and we used cuticular extracts as predator cues (see below for details on the killing and extraction protocol).

##### Experimental setup

Two-patch systems were built inside 30 × 180 cm plastic forcing tunnels (height: 20 cm) placed in a tiled 4 × 4 m room (mean temperature 20 °C; 16L: 8D). Tunnels were duct-taped to the floor in order to prevent escapes. Boxes in which snails were maintained (see above) were placed at one extremity of each tunnel to serve as origin patches; similar boxes, but without snails, were placed on the opposite side to serve as target patches. The between-patch matrix was left empty and dry. With this setup, the between-patch distance was 120 cm; previous observations showed that non-dispersal exploratory movements nearly always spanned distances < 1 m (M. Dahirel, unpublished data).

##### Treatments

One box of ten snails was released per two-patch tunnel, the box itself serving as the origin patch. We tested 5 tunnels per combination of predator cue presence (PRED, yes/no) and food restriction (RA, standard/low) treatments, so 20 tunnels and 200 snails overall (see below for details on the implementation of the treatments). Due to logistical constraints, tunnels were tested in two sessions of ten (start dates: June 9 and 13, 2016); all four possible treatments were roughly equally distributed between these two sessions.

Twenty-four hours before release in the experimental tunnels, the original feeders in each box were removed. They were replaced by new Petri dishes, surrounded with a 3 cm wide ring of Whatman paper with or without predator cues depending on the treatment (see below). Snail food was also added to the new feeders based on treatment (see below). After these 24 hours of habituation, the origin patch box was placed in a randomly selected tunnel, opened, and snails were left free to disperse for 3 days. Whatman papers were left in place, feeders in the origin patch were refilled with food for 3 days (actual quantity depended on treatment), and feeders in the target patch were filled with enough food to sustain all snails for 3 days.

##### Resource availability

During dispersal trials, snails were fed the same cereal flour they were exposed to during maintenance. Previous tests (Dahirel *et al.*, 2016) and new preliminary experiments showed that snails consume on average 0.4 g of flour per day when alone and provided with ad libitum food, independently of snail size. Origin patches in the standard food treatment (standard RA), and target patches in all treatments, were provided with 8 g of snail food per day, i.e. twice the quantity needed to sustain 10 snails based on the above result. It is difficult to evaluate the effect of a short-term food restriction on *Cornu aspersum*, as this species readily enters dormancy; we therefore used the same food restriction as for the land slug *Deroceras reticulatum*, namely −90%. Origin patches in the low resources treatments (low RA) were thus provided with 0.4 g of snail food per day, i.e. 10% of the quantity consumed by 10 snails.

##### Predator cues

*Cornu aspersum*, like most land gastropods, has poor vision and audition, and relies primarily on olfaction to apprehend its environment (Chase, 2001). Cuticular extracts of the predator *Lampyris noctiluca* were extracted following Bursztyka *et al.* (2016)’s protocol. Briefly, beetle larvae that had been starved for 48 h to remove digestive residues were placed in a refrigerator (4 °C) for 4 h. This aimed to induce torpor, avoiding the release of defensive secretions during freezing. Larvae were then transferred to a freezer (−20 °C) for 24 h. Dead frozen beetles were then placed in a new glass vial filled with 20 mL of pure ethanol per gram of live weight. This solution was stored at 4 °C and gently shaken twice a week. This solution was used two weeks after its creation. Pure ethanol stored under the same conditions was used as a control solution. These solutions were then presented to snails using pieces of Whatman paper, with 5 µL of solution per cm^2^ of paper, after leaving paper strips at room temperature for 15 minutes to allow for evaporation of the excess ethanol. To confirm the validity of this extraction protocol and the efficiency of the predatory cue, we placed 12 *Cornu aspersum* snails in individual 9 × 6 × 5 cm boxes with the bottom lined with two strips of Whatman paper, one side with the predatory cues solution, one side with the control solution. After one hour, all snails had chosen a side, and 83.3% (10 out of 12) were on the “control” side (χ^2^ = 5.33, *p* = 0.02).

##### Data collection

The number of dispersing (i.e., in the target patch) and non-dispersing (in the origin patch) snails in each tunnel was recorded once after 3 days.

#### *Cryptomonas* sp

**Authors** Jonathan De Raedt and Frederik De Laender

##### Study organism

The subject of this study was strain 26.80 of *Cryptomonas* sp., acquired from the EPSAG Culture Collection of Algae. The strain was cultured in 500 mL Erlenmeyer flasks closed with semi-permeable lids. Cultures were maintained in COMBO medium (Kilham *et al.*, 1998). Every week, 30 mL of fluid was removed from the flasks and placed in new Erlenmeyer flasks in a 10:1 medium-algae ratio. Algal cultures were maintained in an 16L:8D photoperiod at 19 °C. Because we clonally propagated a single strain of *Cryptomonas* sp., there is essentially no genetic variation among our treatments.

*Daphnia magna* was used as the predator species and cultured in 600 mL beakers filled with COMBO medium at 19 °C. Medium was replenished twice a week. Simultaneously, 5 mL of a 2 to 3 week old *Cryptomonas* sp. culture was added as a food resource.

*Crypotmonas* sp. and *Daphnia magna* are very abundant in most waters of Western Europe. *Daphnia pulex* grazes on phytoplankton, algae and bacteria. The amount of studies investigating effects of predator cues of *Daphnia* is limited. However, it has been shown that the distribution of phytoplankton in naturally occurring freshwater varies in response to the resident zooplankton community (Arvola *et al.*, 1992). Moreover, Latta *et al.* (2009) found that predator cues induced phototactic movement of *Chlamydomonas reinhardtii* that resulted in a higher algae density at the surface than in untreated water. By remaining at the water surface, the algae could avoid its grazer since the grazers avoid the highest water layers in the presence of predators.

Another algae, *H. akashiwo*, was reported to have higher swimming speeds and vertical velocities in the presence of a ciliate predator. The direction was upward in the presence of a halocline. The ciliate was not able to persist at low salt concentrations and, as such, this area was a safe zone for the algae. In the absence of a halocline, the direction of the algae was downward, while the ciliates were aggregated at the top of the tank (Harvey & Menden-Deuer, 2012).

##### Experimental setup

We construct a 2-patch system consisting of 2 1.5 mL Eppendorf tubes connected by a Fluoroelastomer tube (length 2.5 cm, inner diameter 4 mm, outer diameter 6 mm). Fluoroelastomer is black and nontransparent and thus avoids light penetration in the connection tube. As a result, this part of the set-up is considered as the hostile matrix.

##### Treatments

First, 2.78 mL of a PO_4_^3−^ -poor medium was added to all dispersal systems, independent of the treatment. After filling the tubes, the connection tube was closed.

For the control treatment (standard RA, no PRED), 11.2 µL of a KH_2_PO_4_^3−^ -solution was added to both patches to obtain a PO_4_^3−^ -rich medium. Finally, 140 µL algae and 140 µL medium were added to the start and target patch respectively.

For the predator cues treatment (yes PRED), 0.695 mL of medium was replaced with 0.695 mL of a predator cues PO_4_^3−^ -poor medium in the start patch. Next, 11.2 mL of the KH_2_PO_4_^3−^ -solution was added to both patches to obtain a PO_4_^3−^ -rich medium. Finally, 140 µL algae and 140 µL medium was added to the start and target patch respectively.

For the low resource treatment (low RA), 11.2 µL Milli-Q water and 140 µL algae were added to the start patch to maintain a PO_4_^3−^ -poor medium. Next, 11.2 µL of the KH_2_PO_4_^3−^ -solution was added to the target patch to obtain a PO_4_^3−^ -rich medium. Finally, 140 µL medium was added to the target patch.

For the combined predation and low resource treatment (low RA, yes PRED), 0.695 mL of medium was replaced with 0.695 mL of the predator cues PO_4_^3−^ -poor medium in the start patch. Next, 11.2 µL of Milli-Q water and 140 µL algae were added to the start patch. Afterwards, 11.2 µL of the KH_2_PO_4_^3−^ -solution was added to the target patch to obtain a PO_4_^3−^ -rich medium. Finally, 140 µL medium was added to the target patch.

The experiment was run for 18 h (approx. 5% dispersal in control patch). The acclimation phase was 4.5 h (that is, 25% of the duration of the experiment).

There were 6 replicates per treatment. The whole experiment was run in 2 blocks, each consisting of 3 replicates. The blocks differed in incubator and the moment the experiment was started (1 h).

##### Resource availability

Resource: Combo medium (Kilham *et al.*, 1998), with phosphate as limiting resource in the origin patch (low RA). Phosphate is added as KH_2_PO_4_^3−^ (Normal [KH_2_PO_4_^3−^] = 8.71 mg L^−1^; low [KH_2_PO_4_^3−^] = 0.871 mg L^−1^).

##### Predator cues

*Daphnia* is a common predator of *Cryptophyta*. Grazer kairomone water was created by isolating 40 individual *Daphnia* adults and placing them in 100 mL of filtered Combo medium (Kilham *et al.*, 1998) (PO_4_^3−^ -limited: 10%) for 18 h. A similar procedure was followed by Latta *et al.* (2009). Prior to use in microcosms the kairomone water was filtered through 165 µm nitex mesh to remove particulate matter and the *Daphnia*.

##### Data collection

Sampling regime: Both patches were sampled after the dispersal phase. 1.2 mL per patch was removed and stored in 1.5 mL Eppendorf tubes. To preserve the samples and fixate the algae, lugol was added. Densities were determined using a 1 mL counting chamber.

Moreover, a sample of 8 µL from the start patch was taken for video analysis (concentrations were too low to take samples of the target patch for video analysis).

For measuring dispersal, cell numbers were acquired using a counting chamber.

#### Grey field slug — *Deroceras reticulatum*

**Authors** Maxime Dahirel, Armelle Ansart and Luc Madec

##### Study organism and predator

The grey field slug, *Deroceras reticulatum* (Müller; Gastropoda, family Agriolimacidae) is one of the most common and economically important crop pests of temperate regions (Barker, 2002), and is occasionally encountered in forests (Kappes *et al.*, 2009). Slugs used in this experiment were caught using non-baited traps (De SangosseⓇ, Pont-du-Casse, France) in the spring of 2016 in a wheat field in Menetou-Salon, France (47°16^*′*^1^*″*^ N, 2°22^*′*^23^*″*^ E). Slugs were housed in transparent polyethylene boxes (26.5 × 13.5 × 8.5 cm, approx. 50 randomly assigned slugs per box) lined with synthetic foam kept saturated in water. Egg carton pieces saturated in water were added to be used as shelters. All rearing and experimental boxes were kept under controlled conditions (10 *±* 1 °C, 12L: 12D; Armsworth *et al.* 2005). Slugs were fed ad libitum with commercial snail food (cereal flour supplemented with calcium, HélinoveⓇ, Le Boupère, France), cucumber (*Cucumis sativus* Linnaeus) and lettuce (*Lactuca sativa* Linnaeus). Boxes were cleaned, and the shelters and lining changed, twice a week.

The parallel-sided ground beetle *Abax parallelepipedus* (Piller & Mitterpacher; Coleoptera, family Carabidae) was used as our model predator. This large generalist predatory beetle is present in forests and agricultural landscapes in western Europe, although it is rarer in the latter (Duflot *et al.*, 2016; Eyre *et al.*, 2016). Large generalist carabid beetles such as *A. parallelepipedus* are predators of slugs, including *Deroceras reticulatum* (Symondson, 1993; Symondson & Liddell, 1993; Symondson, 2004). Slugs have both much smaller home ranges (in the range of 1 m^2^ versus 14 to 650 m^2^ depending on environments, South 1965, Charrier *et al.* 1997) and slower movement speeds than *Abax parallelepipedus* (maximal speed approx. 10 cm min^−1^ versus up to 20 cm s^−1^, M. Dahirel, personal observations on tested slugs; Forsythe 1981). *D. reticulatum* has previously been shown to be able to detect and avoid several predatory ground beetle species based on various olfactory cues (Armsworth *et al.*, 2005; Bursztyka *et al.*, 2013, 2016). Beetles were caught by hand in April 2016 in a small forest near Pléguien, France (48°36^*′*^ N, 2°56^*′*^ W). They were then maintained in controlled conditions (20 °C *±* 1 °C; 16L: 8D), in 5.5 cm high cylindrical boxes (diameter: 9 cm) with 0.5 cm of moistened soil at the bottom (one to two beetles per box). Beetles were fed ad libitum with moistened dried cat food and apple slices; in addition, one live *Deroceras reticulatum* slug was provided per beetle once a week.

##### Experimental setup

Two-patch systems were built in 40 × 13 cm transparent plastic boxes (height: 9 cm). Boxes were divided in two 10 × 13 cm patches and one central 20 × 13 cm matrix, separated by plastified cardboard walls. Two 13 × 1 cm slots were left open in each cardboard wall to allow slugs to leave and enter patches. One feeder (plastic bottle cap) and one shelter (wet egg carton piece) were present in all patches, in addition to 1 cm of humid soil at the bottom. The between-patch space was left empty and dry. In preliminary tests (ad libitum food, no predator cues; see below), this setup resulted in between 20 and 30% of dispersers after 3 days, which is roughly the time most slugs stopped dispersing away from release in a release-recapture study (South, 1965).

##### Treatments

Ten randomly chosen slugs were released per two-patch tunnel, in one of the patches (hereafter the ‘origin’ patch’); this density (0.13 per m^2^) is at the low end of the wide range of naturally observed population densities (e.g. Symondson *et al.*, 1996; Kappes *et al.*, 2009), as higher densities are difficult to manage in controlled conditions. We only used slugs with fresh masses over 200 mg prior to the experiments, to ensure all individuals used were sexually mature/ reaching sexual maturity (Armsworth *et al.*, 2005). We tested 5 tunnels per combination of predator cue presence (yes/no PRED) and food restriction (standard/low RA) treatments, so 20 tunnels and 200 slugs overall (see below for details on the implementation of the treatments). Due to logistical constraints, tunnels were tested in two sessions of ten (start dates: May 5 and 19, 2016); all four possible treatments were roughly equally distributed between these two sessions. Slugs were placed in the origin patch, which was then closed for 24 h using an upside-down box of the same size. Whatman papers, with or without predator cues depending on the treatment (see below), was placed at the same time in shelters (7 × 7 cm) and around feeders (1.5 cm wide strip) in the origin patch, so that slugs had to crawl on them on their way to eat or rest. Lettuce was also added to feeders based on treatment and slug body mass (see below). After these 24 hours of habituation, the origin patch was opened and slugs left free to disperse for 3 days. Whatman papers were left in place, feeders in the origin patch were refilled with food for 3 days (actual quantity depended on treatment), and feeders in the target patch were filled with enough lettuce to sustain all slugs for 3 days.

##### Resource availability

Food consumption of *Deroceras reticulatum* under optimal availability conditions was estimated by placing slug pairs of known fresh mass for 24 h in 5.5 cm high cylindric boxes (diameter: 9 cm) with 16 cm^2^ of lettuce per box (N = 10 pairs of slugs). Slugs consumed on average 10.02 × slug mass (g) + 0.08 cm^2^ of lettuce per day per slug (*R*^2^ = 0.71). Slugs in the standard RA treatments were then provided with the quantity they were expected to consume during the experiment plus 24 h of surplus, based on the above linear equation. Slugs in the low RA treatments were provided with 10% of the expected quantity; preliminary observations showed that −90% was the strongest restriction that did not result in outright cannibalism attempts, as opposed to mere agonistic interactions, within the duration of the experiment. Food consumption was measured as the difference between the lettuce leaf surface present at the end of the 24 h-habituation period and the surface present at the end of the 3-day dispersal test.

##### Predator cues

*Deroceras reticulatum*, like most land gastropods, has poor vision and audition, and relies primarily on chemical cues to apprehend its environment (Chase, 2001). Predator cues isolation was inspired by Armsworth *et al.* (2005). Ten *A. parallelepipedus* beetles were left to walk one night (10 h) in an otherwise empty 22 × 17.5 × 9.5 cm box lined with Whatman paper strips (total surface: 420 cm^2^ of paper per box). Pristine Whatman paper strips were used as controls. Before being used in our experiments, this cue isolation method was tested following Bursztyka *et al.* (2013). Briefly, 26.5 × 13.5 × 8.5 cm test boxes were divided in two darkened shelters with a small 1 cm entrance and one central lit part. Wet egg cartons pieces and a feeder with commercial snail food were present in both shelters. In each test box, one randomly chosen shelter had a 1.5 cm wide Whatman paper strip with predator cues placed at its entrance; the other had a control paper strip. Slugs were placed, one at a time, in the lit central part of their box and were left free to move for 24 h. After this time, most slugs were found in the control shelter instead of the one with the predator cue (90%; 18 out of 20; *χ*^2^ = 12.8; *p* < 0.001). Four other carabid taxa were tested at the same time using the same protocol and sample size (*Poecilus cupreus* (Linnaeus); *Nebria brevicollis* (Fabricius); mixture of *Pterostichus madidus* (Fabricius) and *P. melanarius* (Illiger)); *A. parallelepipedus* was the one eliciting the strongest response, and the only one in which all individual beetles ate all provided live slugs within 24 h.

##### Data collection

The number of dispersers (i.e., individuals in the target patch) and residents (individuals in the origin patch) in each tunnel was recorded once after 3 days.

#### Amphipods — *Dikerogammarus villosus* and *Gammarus fossarum*

**Authors** Chelsea J. Little, Emanuel A. Fronhofer and Florian Altermatt

##### Study organisms and predator

We used two amphipod (Crustacea, Amphipoda) species as study organisms: *Gammarus fossarum* (Koch) and *Dikerogammarus villosus* (Sowinsky). Amphipods are key macroinvertebrates in stream, river, and lake food webs because of their dual roles in shredding terrestrial detritus and serving as food items for larger organisms. Amphipods show a range of responses to fish predation, including reduction of drifting behavior, drifting primarily at night-time, finding refuge in benthic macrophytes, remaining in sediments, or simply sitting motionless to avoid visual detection; by contrast, the presence of macroinvertebrate predators can increase drifting behavior (Macneil *et al.*, 1999). In general, habitat complexity provides a respite from fish predation (Diehl, 1993). Amphipods have two main modes of dispersal: a more active swimming or crawling mode, where 25-33% of individuals disperse (Hughes, 1970; Elliott, 2003), and a less-active drifting mode of dispersal. Because drift is not relevant in our experimental setup since there is no current (see below), we focus on the more active form of dispersal.

*G. fossarum* is a small freshwater amphipod native and common to central Europe. In November 2016 we collected individuals by kicknet from the third-order Sagentobelbach stream in Dübendorf, Switzerland (47.39° N, 8.59° W). We selected adults in medium and large size classes, and later distributed them into experimental units such that distributions of size classes was uniform across the experiment. It is impractical to identify individuals by sex in the field/while living, except by separating precopulatory pairs; perhaps partially due to the season, we found only a few such pairs, and thus collected individuals without regard to sex. We assume that allocation of individuals to treatments was relatively even across the experiment. Amphipods were brought to the lab and placed in large holding containers of approx. 500 individuals, where they were gradually brought up to 18 °C, fed alder (*Alnus glutinosa* (Gaertner)) leaves (which had been conditioned in stream water with natural microbial and fungal communities for six days) ad libitum, and maintained for two and a half days before being allocated to experimental units.

*D. villosus* is a larger freshwater amphipod native to the Ponto-Caspian region which has established itself through the Rhine catchment in the last two decades (van den Brink *et al.*, 1991). In January 2017 we collected individuals by kicknet from Lake Constance at Kesswil, Switzerland (47.60°N, 9.32° W). Collection with respect to size and sex, and maintenance in the laboratory before the experiment, was identical as for *G. fossarum*.

The European perch (*Perca fluviatilis* (Linnaeus)) was used as a predator for both amphipod species. Perch is a highly mobile predator in lakes, and feeds on both other fish and on macroinvertebrates. Gammarid amphipods can make up a large part of the diet of lake perch (Allen, 1935; Persson, 1981; Jamet & Desmolles, 1994), and young adult perch can provide strong top-down control of macroinvertebrate biomass (Diehl, 1993). In the lower Rhine, the diet of perch has shifted substantially with the arrival of non-native species, and includes more non-native amphipods (Kelleher *et al.*, 1998). We obtained fresh-caught, dead perch (Fischerei Grieser, Obermeilen, Switzerland for experiments with *G. fossarum* and Braschler’s Comestibles, Zurich, Switzerland for experiments with *D. villosus*) and immediately swabbed the sides of the fish with cotton balls to capture mucus and the chemical cues contained therein (see below for details).

##### Experimental setup

Two-patch systems were built using 3 L (198 × 198 mm) polypropelene boxes connected by approx. 30 cm of silicon tubing with an inside diameter of 20 mm (previous observations with multiple tubing sizes showed that this diameter led to 17% dispersal of *G. fossarum* from control patches over a five-hour period; Little and Fronhofer, unpublished data). All boxes contained either alder leaves or imitation cloth leaves (see below for details) as well as a plastic imitation macrophyte to provide shelter and habitat complexity. Origin and target patches were randomized with respect to directionality. All boxes/patches were covered with a black lid to reduce light permeability, while the connection tube/matrix was left uncovered. Prior work showed that when large tanks were half-covered with the shading lid, only 20% of *G. fossarum* stayed in the unshaded portion, thus preferring a shaded habitat (Little and Fronhofer, unpublished data).

##### Treatments

At the beginning of an experiment, 20 amphipods were placed in each origin patch. We tested ten replicates per combination of predator cue presence (yes/no PRED) and food restriction (standard/low RA) treatments, so 40 tunnels and 800 amphipods per species (see below for details on the implementation of the treatments). Due to logistical constraints, experiments on the two species were run separately (*G. fossarum* start date: 10 November, 2016; *D. villosus* start date: 26 January, 2017).

Twenty-four hours before amphipods were placed in the experimental units, real or imitation leaves were placed in the origin and target patches (see below). The connection tube was shut using clamps to prevent floating movement of resources between the patches. At the same time as amphipods were placed in the origin patches, cotton balls containing predator cues, or clean control cotton balls, were also placed in the origin patches (see below). After 30 minutes of habituation, the connection tube was opened and amphipods were left free to disperse for 4.5 h (*G. fossarum*) or 7 h (*D. villosus*). At the end of the dispersal phase, the connection tubes between origin and target patches were closed again with clamps and all amphipods were removed from the patches.

##### Resource availability

During dispersal trials, amphipods were fed the same alder leaves they were exposed to during maintenance. Origin patches in the standard RA treatment, and target patches in all treatments, were provided with 1.5 g dry weight of alder leaves. Because amphipods can survive for at least two weeks under starvation conditions (Nyman *et al.*, 2013), we decided to use a complete food restriction and offered only imitation cloth leaves in the restricted-food treatment. The number of imitation leaves were chose to roughly match the surface area provided by the alder leaves in the standard RA treatment, in order to minimize any confounding effects of amphipods using the leaves themselves as habitat or hiding spots (Haeckel *et al.*, 1973). Food consumption was not measured in this experiment because previous experiments in the same laboratory conditions showed that *G. fossarum* consume 0.35 mg dry weight of alder leaves per mg dry weight of amphipod per day, and *D. villosus* 0.5 mg dry weight of alder leaves per mg dry weight of amphipod per day (Little & Altermatt, submitted); with only a few hours of experimental duration, it would be difficult to measure leaf consumption in the experimental units with sufficient precision to detect differences between treatments, if they did exist. 1.5 g dry weight of alder leaves were provided in all target patches.

##### Predator cues

Cotton balls wiped on the sides of freshly killed, commercially caught *P. fluviatilis* to collect their mucus and the contained chemicals were used use as predator cues. Cotton balls were frozen at −20 °C until the day of the experiment. Conspecific amphipods have been shown to alter their activity in the presence of fish infochemicals (Dunn *et al.*, 2008; Schäffer *et al.*, 2013; Szokoli *et al.*, 2015), and a pilot experiment showed that these perch predator cues reduced dispersal rates from 5% to 0% from non-food-limited patches by *G. fossarum*. One cotton ball was placed in each origin patch in the predation treatment (yes PRED) units, and a clean, sterile cotton ball (which had also been frozen at −20 °C) was placed in the origin patch in no-predation treatment (no PRED) units.

##### Data collection

The number of dispersers (i.e. individuals in the target patch) and residents (individuals in the origin patch) in all experimental unit was recorded once after the dispersal period was concluded.

#### Palmate newt — *Lissotriton helveticus*

**Authors** Delphine Legrand, Alexandre Vong, Laurane Winandy and Julien Cote

**License** 09-2016-02 (palmate newt); 2012-10 DREAL (grass snake)

##### Study organism and predator

The palmate newt (*Lissotriton helvetucus*, Razoumovsky, 1789) is a small European newt species (adult length: males, up to 85 mm; females, 95 mm). In this experiment, we used 216 small newts (17.8 *±* 2.6 mm SE) in the terrestrial phase captured in the Metatron (Legrand *et al.*, 2012) and maintained in 4 terrariums (35 × 17.5 × 22.5 cm, 54 individuals per terrariums). Terrariums contained 5 cm of soil litter covered with mosses, 2 pieces of egg carton used as refuges, two small dishes for water and a regular addition of bloodworms. As we caught newts in semi-natural enclosures, they have never been exposed to predator cues (see below).

The grass snakes (*Natrix natrix*) were used as our model predator. Grass snakes are active and generalist feeders, preying upon amphibians, fish, small mammals, reptiles and birds (Hailey & Davies, 1986; Gregory & Isaac, 2004; Consul *et al.*, 2009). Grass snakes forage on newts mainly in their aquatic phase. We used several snakes maintained for another study in a reptile facility room.

We estimated that newts had similar home ranges as snakes (< 50 000 m^2^ for newts, Verell 1987; Hehl-Lange 2001, 12000 – 36 000 m^2^ for grass snakes, Madsen 1984; Reading & Jofré 2009), while the mobility, estimated as sprint speed, was higher for snakes than for newts (newts: < 10 cm s^−1^, in another newt genus, Wilson 2005; Gvoždík & Van Damme 2006; snakes: 30 – 60 cm s^−1^, Hailey & Davies 1986; Isaac & Gregory 2007). On top of dispersal, newts can exhibit a postural defence exposing ventral orange colouration (Brodie, 1977). During this posture there is a release of skin secretion (tetrodotoxin) making them inedible to predators (Yotsu-Yamashita *et al.*, 2007).

##### Experimental setup

We used 8 dispersal systems placed in a greenhouse with controlled temperature (16 – 25 °C). Each system is made of two 130 L plastic containers (78 × 56 × 43 cm) connected by a circuitous plastic pipe (diameter: 10 cm, total length: 4.4 m) on the upper section of the container. The departure container was filled with soil litter providing access to the corridor while we only added a thin layer of soil in corridors and arrival containers. To go from the departure to the arrival containers, newts had to enter this narrow corridor and fall into the arrival container. Before releasing the newts, we took ventral and dorsal pictures to allow an individual identification.

##### Treatments

We created 24 populations made of 8 – 11 newts (approx. 8.23 *±* 0.29 SE). We used a 2 × 2 factorial design, crossing resource availability (RA) and predation risk (PRED) with two levels each, resulting into six replicates of each combination of treatments. We ran the eight replicates in 3 blocks, from November 23rd to December 15th 2016. The RA and PRED treatments were applied before releasing newts for a 24 hours acclimation phase with connections between containers closed. After 24 hours, connections were opened and we monitored dispersal movements as described below.

##### Resource availability

RA included two treatments: a low and high resources treatment. The resources were manipulated through the addition of approx. 90 bloodworms in the high resources treatment and approx. 10 in the low food resources, ensuring a large difference in food availability between high and low food enclosures.

##### Predator cues

Snakes were housed in individual plastic box containing a water dish and a piece of humid absorbent paper to collect odours serving as predator cues. For each population, we used an absorbent paper belonging to different snake boxes. Before releasing newts, dispersal systems with predator cues treatments received a piece of absorbant paper from a snake box, whereas control treatments received a piece of odour free paper. When removing a paper from a snake box, we added a new piece of paper for future experimental blocks. Similar protocol to collect snake olfactory cues has been shown to be efficient in eliciting antipredator reaction in terrestrial salamander species (Madison *et al.*, 1999; Maerz *et al.*, 2001; Sullivan *et al.*, 2002). While newts’ responses to predator chemical cues have been mostly studied in the aquatic phase (Winandy & Denoël, 2013), newts in the terrestrial phase are known to use olfactory cues to select habitats (Joly & Miaud, 1993).

##### Dispersal and data collection

One day after the opening of corridors, we stopped dispersal and captured back newts in the departure and arrival containers. At the end, we recaptured and identified 177 newts which were used in our analysis.

#### Paramecium caudatum

**Authors** Florent Manzi and Oliver Kaltz

##### Study organism and predator

We used *Paramecium caudatum* as focal species (prey) and *Didinium nasutum* as predator. The *Paramecium* were taken from a long-term selection experiment on dispersal (O. Kaltz, unpubl. data; see below); the founder population for this experiment comprised a mix of 20 clones, but at the time of the present experiment, individual selection lines most likely consisted of single clones. Preliminary analysis indicates that different clones are fixed in high-dispersal and low-dispersal selection lines. *Didinium* was obtained from Sciento (strain P220).

This predator-prey system naturally occurs in fresh water environments, with *Didinium* feeding on different species of the genus *Paramecium* (Veilleux, 1979). *Paramecium aurelia* shows predator-induced dispersal in the presence of a flatworm predator (Hammill *et al.*, 2015). Other ciliate species can detect the presence of predators by direct membrane contact (Kuhlmann, 1994) or use the hydrodynamic disturbance induced by cilia motion (Kusch, 1993). In a pilot experiment, we found a tendency of increased dispersal of our *P. caudatum* strains exposed to a filtrate prepared from a *Didinium culture* (F. Manzi, unpubl. data). This suggested a plastic dispersal response mediated by chemical cues.

Both *Paramecium* and *Didinium* cultures were maintained in temperature-controlled incubators, at 23 °C in the dark. For the *Paramecium* cultures, we used an organic lettuce medium (1 g of dried lettuce suspended in 1.5 L of Volvic™ mineral water), supplemented with the bacterium *Serratia marcescens* (Nidelet & Kaltz, 2007). The *Didinium* were regularly (2–3 times per week) fed with a mix of *Paramecium* from the above-mentioned selection experiment.

Similar to Fronhofer & Altermatt (2015), the long-term lines of *Paramecium* were going through alternating cycles of dispersal (3 h) and logistic growth (7 d), for approximately 1 year. For the present experiment, *Paramecium* were taken at the end of a growth cycle, when populations had reached carrying capacity.

We used independent *Paramecium* selection lines from each of two long-term directional selection treatments: high-dispersal lines and low-dispersal lines.

##### Experimental setup

The two patches consisted of 13 mL plastic tubes with a round bottom, connected by silicone tubing (length: 5 cm, diameter: 0.8 mm), allowing dispersal of *Paramecium* between the two patches. The connection can be opened and blocked by means of plastic clamps.

To measure dispersal, *Paramecium* were filled in one of the two tubes (‘patch of origin’), while the connection between the two tubes was blocked. After an acclimation period of 30 min (approx. 17% of the duration of the dispersal phase), the clamps were removed for three hours, during which time the *Paramecium* could freely swim between tubes. Then the clamps were put back to block the connection. ‘Dispersers’ are defined as individuals that are present in the ‘target patch’ at the end of the 3 h period. To estimate the proportion of dispersed individuals (i.e., the dispersal rate), the density of individuals in origin and target patches were estimated immediately after blocking the connections, by by counting the number of individuals in 300 µL samples from the patch of origin and 1 mL samples from the target patch. Replication during the dispersal period can be neglected (Fellous *et al.*, 2010).

This experimental setup corresponds to the dispersal selection conditions of the above-mentioned long-term selection experiment. In high-dispersal selection treatments, non-dispersing *Paramecium* are discarded, and only dispersing *Paramecium* from the target patch are retained, propagated for one week and then subjected to a new round of 3 h dispersal, and so on for 1 year. In the low-dispersal treatment, only the non-dispersing *Paramecium* are retained and used for a new growth/dispersal cycle. The experiment was conducted at 23 °C.

Standard resource (standard RA) conditions consisted of sterilised lettuce medium (90% volume), to which *Serratia* from a stock culture was added as a food resource (10% volume). Prior to use, the medium was filtered through two layers of sterile medical gauze to remove larger lettuce particles.

The connection between the tubes (i.e., the matrix) was filled with sterile mineral water (Volvic™). This was achieved by first filling the entire two-patch system with water, then isolating the matrix from the patches by way of clamping, and replacing the water in the patches by medium and *Paramecium*, according to treatments.

The target patch always contained 13 mL of standard medium. The composition of the patch of origin varied according to treatment (see below).

##### Treatments

We used a total of 12 *Paramecium* selection lines of (6 lines from the ‘high-dispersal’ treatment and 6 lines from the ‘low-dispersal’ treatment) for the experiment. Each line was used at its ‘natural’ density (i.e., density reached after one week of culture) and we did not correct the total number of individuals present in the patch of origin of a given replicate.

Prior to the experiment, two 25 mL samples from each selection line were centrifuged for 30 min at 1500 g, and then approx. 22.5 mL of the supernatant discarded. For one sample, the concentrated *Paramecium* were then resuspended in standard medium to give a new total volume of 15 mL; from this volume the two high-resource replicates were established in the experiment (see below). The other sample was resuspended in sterile mineral water to give two low-resource replicates.

On average, the final number of individuals introduced in the patch or origin was 1093 *±* 96 s.e. There was no significant difference between high-dispersal and low-dispersal lines (*p >* 0.2).

We ran a total of 12 independent replicates for each treatment, giving a total of 48 replicates, with each selection line represented by one replicate in each treatment (2 RA treatments × 2 PRED treatments × 2 selection origins × 6 selection lines). The experiment was organised in six blocks of 8 replicates, with each treatment replicated twice per block. There was a delay of approx. 35 min between the start of each block.

##### Resource availability

The resource consumed by *Paramecium* is the actively hunted bacterium *Serratia marcescens*. This bacterium is motile and may be able to disperse between patches. We did not establish whether this actually happened during the experiment.

To set up standard- and low-resource treatments (standard/low RA), we centrifuged our *Paramecium* cultures (see above) and discarded almost all the supernatant, thereby removing (unsedimented) *Serratia*. In the standard-resource treatment, the patch of origin contains 6 mL of *Paramecium* (previously centrifuged, retrieved from the pellet and diluted in fresh culture medium) and 3.5 mL of either predator filtrate or control filtrate. The volume is completed to 13 mL using 3.5 mL of ‘high food’ medium.

In the low RA treatment, the patch of origin contains 6 mL of *Paramecium* (previously centrifuged, retrieved from the pellet and diluted in sterilized water) and 3.5 mL of either predator filtrate or control filtrate. The volume is completed to 13 mL using 3.5 mL of sterilized water.

We did not quantify the density of *Serratia* in our resource treatments. Due to our dilution protocol, the *Serratia* content in our the low-resource treatment was reduced by c. 70% relative to the high-resource treatment (which corresponds to ad libitum conditions). In a growth assay prior to the main experiment, the low-resource treatment reduced *Paramecium* population growth rate by 80% (data not shown).

##### Predator cues

Other ciliate species (e.g., *Euplotes octocarinatus*) detect the presence of predator by direct membrane contact or use the hydrodynamic disturbances induced by cilia motion (Kusch, 1993; Kuhlmann, 1994). Here, we make the assumption that *P. caudatum* detect the presence of *Didinium* through the use of chemical cues.

We combined 10 mL from each of 10 populations containing both *Paramecium* and *Didinium* at variable densities (overall, c. 5 × 103 *Paramecium* and *Didinium*, respectively). These cultures had been set up several weeks prior to the experiment, and were regularly supplied by a mix of *Paramecium* from high- and low-dispersal selection lines to maintain the *Didinium* populations. The combined populations were centrifuged at 1500 g for 30 min. The supernatant was retrieved and filtered through a micropore filter (0.2 µm). This filtrate served as *Didinium* cue (‘predator filtrate’; yes PRED).

In the same way, we prepared the control filtrate (no PRED). Here, our lettuce culture medium served as a basis, without any addition of *Serratia*, *Paramecium* or *Didinium*. Prior tests had shown that filtrates from pure medium and filtrates from *Paramecium* cultures (without *Didinium*) did not significantly differ in the effect on dispersal; and the effect of both types of filtrate were not significantly different from a sterile water control (results not shown).

A pilot experiment (data not shown) indicated a positive relationship between dispersal rate and filtrate concentration (0% – 10% – 100%), as well as the number of *Didinium* individuals added (0 – 5 – 25).

##### Dispersal and data collection

*Paramecium* were allowed to disperse for 3 h, until the connection between origin and target tubes was blocked. We then took 300 µL samples from the tube of origin and 1 mL samples from the target tube. The number of individuals in these samples were counted under a dissection microscope (20x). From these measurement we extrapolated the total number of individuals in both tubes in a total of 13 mL.

The entire experiment was conducted within one day, with maximal intervals of 30 min between blocks.

Density measurements were taken immediately after the 3 h of dispersal, in the tube of origin and in the target tube. From these measurements we calculated total population size and the proportion of dispersal. As mentioned above, each selection line was used at its ‘natural’ density, i.e., density reached after one week of culture.

#### European minnow — *Phoxinus phoxinus*

**Author** Simon Blanchet

**License** E-2016-130

##### Study organism and predator

The European minnow (*Phoxinus phoxinus*) is a small bodied-size (max. length of approx. 8 cm) freshwater fish species widespread in Europe. Its main predators include predatory fish as well as piscivorous birds and mammals. We here used 192 adults (3-years, approx. 6.2 cm long, min: 4.5 cm max: 8.4 cm) fish originating from a single population raised in artificial lakes in Brittany (France). Minnows were reared for two weeks in a 1100 L external tank and feed ad libitum with pellets and frozen bloodworms.

We used chemical cues from a common predatory fish (the Eurasian perch, *Perca fluviatilis*) that is widely co-occurring with minnows throughout their ranges. A single perch (approx. 20 cm in length) was captured and maintained alive in a 800 L external tank.

We estimated minnow and perch home ranges from stationary distance obtained from the ‘fishmove’ R package (Radinger & Wolter, 2014) and that we considered as the home range radius. We estimated this stationary distance for 30 days using a species and size specific approach for the perch (length = 200 mm) and a size specific approach for minnows (length = 80 mm), because species information was unavailable. Minnows have a smaller home range than the perch (80 m^2^ for minnows, 5808 m^2^ for Eurasian perch). The mobility, estimated as sprint speed, was also smaller for minnows than for the perch (minnow: approx. 40 cm s^−1^, estimated on close species from Mee *et al.* 2011; 183 cm s^−1^ for other perch species, Mallen-Cooper 1994). On top of dispersal, minnows use schooling as an efficient antipredator defence (Magurran & Girling, 1986).

##### Experimental setup

The experimental setup consisted in a stepping stone system made three circular 1100 L plastic tanks connected with opaque pipes of 16 cm in diameter and 120 cm in length. Each of the three tanks was filled with tap water at least 4 days before the experiment started. The bottom of each tank was not filled with medium in any of the tanks. At the end of the last pipe, we added a cut plastic bottle to create an anti-reverse device and prevent dispersers to return in departure tanks.

##### Treatments

We created 24 populations made of 8 fish that were from the same cohort and from a single strain to avoid age and strain effects. Individuals were not sexed before or after the experiment. We used a 2 × 2 factorial design, crossing resource availability (RA) and predation risk (PRED) with two levels each. We ran the eight replicates in 2 blocks (November 3rd to 10th 2016 and December 2nd to 9th 2016). Three days before the experiment started, groups of eight minnows were encaged in 5 L bottles directly emerged in their rearing tank. The fish were then acclimatized for 24 hours with RA and PRED treatments in the departure tank before opening the corridors. After 24 hours, connections were opened and we monitored dispersal movements as described below.

##### Resource availability

RA includes two treatments a low and high resources treatment. The resources are manipulated through the addition of frozen bloodworms (approx. 8 g) in the high resources treatment whereas we did not introduce food in the ‘low food’ treatment. It allows creating a large difference in food availability between high and low food enclosures.

##### Predator cues

Minnows have already been shown to behaviourally react to the odour of their natural predators (Murphy & Pitcher, 1997), although these reactions were exacerbated when predatory odours were mixed with Schreckstoff (Magurran 1989; Blanchet S., unpublished data). To maximize the potential reaction of minnows, we therefore mixed these two types of alarm in our experiment. Specifically, for each tank with predator cues, 200 mL of water was extracted from the perch tank and mixed (using a grinder) with the skins of two freshly dead minnows (Magurran, 1989). We used this solution to mimic the presence of a dangerous predator in our experiment.

##### Dispersal and data collection

We allowed fish to disperse for 7 days during the first block and 10 days during the second block. All tanks were then emptied out to count the number of fish in each tank. We considered the residents as the fish having been caught in the departure tank, while dispersers were those in the farthest tank from departure tanks. It resulted into 125 fish used in the analyses.

#### Large white butterfly — *Pieris brassicae*

**Authors** Delphine Legrand, Staffan Jacob and Julien Cote

**License** 09-2016-02

##### Study organism and predator

The large white butterfly (*Pieris brassicae*) is a widespread butterfly across Europe (Feltwell, 1982). We used 88 butterflies issued from a breeding (2 generations of breeding initiated in the Metatron (Legrand *et al.*, 2012) from clutches collected in 2016 in Ariège, France). Individuals were all measured for several phenotypic traits (see below), sexed and marked with a specific number on their wings.

The common toad (*Bufo bufo*) was used as our model predator. The species inhabits wet locations in Europe and is a common predator of many insects including adult butterflies (Crnobrnja-Isailović *et al.*, 2012). Eight toads were caught by hand near the Metatron and were maintained as described below.

We estimated that butterflies have a larger home range than the common toad (8000 – 28 000 m^2^ for common toads, Parker & Gittins 1979, *>* 300 000 m^2^ for butterflies, Jones *et al.* 1980, Shreeve 1981). Mobility, estimated as flight speed for butterflies and sprint speed for toads, was higher for butterflies than for toads (butterflies: approx. 500 cm s^−1^ for Pieridae species in natural conditions Srygley & Dudley 1993, and approx. 3 m s^−1^ for *P. brassicae* in experimental conditions, S. Ducatez pers. comm.; toad: approx. 30 m s^−1^, Beck & Congdon 2000).

##### Experimental setup

We used the Metatron, a unique experimental platform dedicated to the study of dispersal in terrestrial organisms that allows the manipulation of both spatial and climatic effects (Legrand *et al.*, 2012), in particular in the large white butterfly (Legrand *et al.*, 2015). We used 16 cages of the Metatron (each 200 m^3^, 10 × 10 × 2 m and covered with insect-proof nets) to create eight pairs of two connected patches. We connected each departure cage (in which butterflies were released) to an arrival cage using a corridor. Thus, butterflies could either remain in the departure cage or freely cross a corridor to join the arrival cage, with the possibility of returns to the departure cage. However, the narrow, S-shaped 19 m long corridors were particularly challenging to cross (i.e., dark and warm conditions, low vegetation) and allow discrimination between disperser and resident individuals (Legrand *et al.*, 2012). We maintained the vegetation of higher height in all departures and arrival enclosures, and added a water pond (25 L plastic container, 60 × 39 × 16 cm). We have previously shown that these conditions allowed the discrimination of dispersal events in *P. brassicae* (Trochet *et al.*, 2013; Legrand *et al.*, 2015). In this study, we also added a small fenced water pond in all departure enclosures for the predation treatment.

##### Treatments

Eleven butterflies (55% females) were released in each departure enclosure which represents a density at the lower range of naturally observed populations (5 – 800 individuals per 100 m^2^, Shapiro 1970). After releasing butterflies, we applied the food resources and the predator cues treatments. We used a 2 × 2 factorial design, crossing resource availability (RA) and predation risk (PRED) with two levels each, resulting into 2 departure enclosures of each combination of treatments. The RA and PRED treatment were applied at the beginning of the acclimation phase. During the acclimation phase, which lasted 24 hours, corridors were kept closed to prevent any movements between departure and arrival enclosures. After 24 hours, corridors were opened and we monitored dispersal movements as described below.

##### Resource availability

RA includes two treatments: a low and high resource treatment. We maintained the vegetation of low height in all departures and arrival enclosures and, in the high food enclosures only, we added 2 feeding flowerpots placed in the same corner, and host plant (fresh cabbages) in the center of all enclosures. This addition allows creating a difference in food availability between high and low food enclosures.

##### Predator cues

PRED includes a predation risk treatment, where two toads were added in the fenced pond, and a no predation risk where the fenced pond was left empty. The addition of toads produced both a visual and olfactory cue with no actual predation. We removed most of other potential predators (i.e., other amphibians, reptiles, spiders) before the experiment. We also added two individuals that were crushed in a tube in each predator treatment as alarm cues of predation, and an empty tube in no predation treatments.

##### Dispersal and data collection

After the opening of corridors, we monitored dispersal daily for 4 days. To do so, two observers entered and walked quietly for 5 minutes in each cages. When a butterfly was located, the observer recorded the individual number. This procedure is commonly used on this species (Legrand *et al.*, 2012; Trochet *et al.*, 2013; Legrand *et al.*, 2015) and we did not observe any individuals leaving the cage during dispersal monitoring. We attributed a disperser status to individuals which moved at least once between cages and a resident status to individuals which never left the departure cage. We retained individuals which were alive after three days and excluded individuals which were never observed as their dispersal status is unknown.

#### Pirata latitans

**Authors** Delphine Legrand, Alexandre Vong and Julien Cote

**License** 2012-10 DREAL

##### Study organism and predator

The wolf spider *Pirata latitans* (Blackwall, 1841) is a European species of the family Lycosidae (body length: 3.5 – 5 mm) occurring in wet and marshy areas. Wolf spiders are generalist and active hunters. In this experiment, we used 257 spider captured in the Metatron (Legrand *et al.*, 2012) and maintained in small tanks (5 L, 28 × 20 × 14 cm). Tanks contained 5 cm of soil litter, tiles used as refuges, a regular addition of cricket (*Acheta domestica*), and were sprayed with water twice a day. As we caught spiders in semi-natural enclosures, they have never exposed to predator cues (see below).

The common lizard (*Zootoca vivipara*) was used our model predator. Common lizards are generalist and active feeders preying upon various arthropods species with a noticeable preference for spiders (Avery, 1966). Lizards were raised in cattle tanks as described in the common lizard methods section (see below).

We estimated the prey and predator species to have home ranges of similar magnitudes (300 – 900 m^2^ for other wolf spiders, Framenau 2005, < 1200 m^2^ for common lizards, Clobert *et al.* 1994). The mobility, estimated as sprint speed, was also similar (lizards: 30 – 60 cm s^−1^, Van Damme *et al.* 1991; Sorci *et al.* 1995, 20 cm s^−1^ for spiders of the same genus, Apontes & Brown 2005).

##### Experimental setup

We used 8 dispersal systems placed in a greenhouse with controlled temperature (16 – 25 °C). Each system is made of two 130 L plastic containers (78 × 56 × 43 cm) connected by a circuitous plastic pipe (diameter: 10 cm, total length: 4.4 m) on the upper section of the container. The departure container was filled with soil litter providing access to the corridor while we only added a thin layer of soil in corridors and arrival containers. To go from the departure to the arrival containers, spiders had to enter this narrow corridor and fall into the arrival container.

##### Treatments

We created 24 populations made of 8 – 10 spiders (approx. 9.83 *±* 0.10 SE). We used a 2 × 2 factorial design, crossing resource availability (RA) and predation risk (PRED) with two levels each, resulting into 6 replicates of each combination of treatments. We ran the 6 replicates in 3 blocks, from July 29h to September 6th 2016. The RA and PRED treatments were applied before releasing spiders for a 24 hours acclimation phase with connections between containers closed. After 24 hours, connections were opened and we monitored dispersal movements as described below.

##### Resource availability

RA includes two treatments a low and high resources treatment. The resources were manipulated through the addition of 100 crickets in the high resources treatment and 25 in the low food resources. It allows creating a large difference in food availability between high and low food enclosures.

##### Predator cues

Lizards were housed in individual terraria containing 3 cm of soil, a shelter (a piece of eggs carton), a water dish, and a piece of absorbent paper to collect odours serving as predator cues. In one corner of the terrarium, ultraviolet and incandescent lamps provided light and heat for thermoregulation from 9:00 to 12:00 and from 14:00 to 17:00. Lizards were fed daily with 1 cricket (*Acheta domestica*). Before releasing spiders, dispersal systems with predator cues treatments received a piece of lizard paper mixes and a piece of egg box, whereas control treatments received a piece of odour free paper and egg box collected from vacant terraria maintained under the same conditions than inhabited terraria. When removing a paper from lizard or lizard-free terraria, we added a new piece of paper for future experimental blocks. The protocol to collect lizard olfactory cues is a standard protocol already used several times to elicit social reactions in lizards (e.g., Cote & Clobert, 2007; Teyssier *et al.*, 2014).

##### Data collection

After the opening of corridors, we monitored dispersal daily for 15 days. To do so, we captured spiders in the arrival containers without disturbing the departure containers. At the end of the dispersal assay, we captured all spiders in departure or arrival container, as well as in the corridor. We were able to recapture only 153 spiders. We only used recaptured ones in our analysis to prevent confounding dispersal with the probability of recapture.

#### White-legged damselfly — *Platycnemis pennipes*

**Authors** Delphine Legrand, Lieven Therry, Alexandre Vong and Julien Cote

**License** 09-2016-02

##### Study organism and predator

The white-legged damselfly (*Platycnemis pennipes*) is a widespread damselfly occurring from the Atlantic to the Jenisei river in Siberia and typically breeds in a wide range of aquatic habitats such as rivers, canals and fish ponds (Dijkstra & Lewington, 2006). We caught 160 damselflies near the Metatron (Legrand *et al.*, 2012) in Ariège (France) in mid-July 2016. Individuals were all measured for several phenotypic traits, sexed and marked with a unique number on their wings. After marking all of them, individuals were transferred to 8 semi-natural enclosures randomly, except for sexes which were distributed to obtain similar sex-ratio among cages. Two hours after releasing damselflies and before applying treatments, we recorded their flying behaviour (see below).

The common frogs (*Rana temporaria*) were used as our model predator. The species inhabits wet locations in Europe and is a common predator of adult damselflies (Corbet, 1999). Eight Frogs were caught by hand in the same location as damselflies and were maintained as described below.

We estimated that damselflies have a larger home range than the common frog (12 500 m^2^ for damselfly, Bennett & Mill 1995, 330 – 1500 m^2^ for common frogs, 330 m^2^: Loman 1994, 1500 m^2^: Haapanen 1970). Mobility, estimated as maximal flight speed for butterfly and sprint speed for frogs, was higher for damselflies than for frogs (Frogs: 20 – 80 cm s^−1^, for another *Ran* species, Miller *et al.* 1993, 140 cm s^−1^, Rüppell 1989). On top of dispersal, damselflies (including *Platycnemis* sp.) exhibit group oviposition to reduce predation risk at the oviposition site where frogs are encountered (Rehfeldt, 1990; Martens, 2002).

##### Experimental setup

We used the Metatron, a unique experimental platform dedicated to the study of dispersal in terrestrial organisms that allows the manipulation of both spatial and climatic effects (Legrand *et al.*, 2012), especially in flying insects (Legrand *et al.*, 2015). We used 16 cages of the Metatron (each 200 m^3^, 10 × 10 × 2 m and covered with insect-proof nets) to create eight pairs of two connected patches. We connected each departure cage (in which damselflies were released) to an arrival cage using a corridor. Thus, damselflies could either remain in the departure cage or freely cross a corridor to join the arrival cage, with the possibility of returns to the departure cage. However, the narrow, S-shaped 19 m long corridors were particularly challenging to cross (i.e., dark and warm conditions and low vegetation) and allow discrimination between dispersal and resident individuals (Legrand *et al.*, 2012). We maintained the vegetation of higher height in all departures and arrival enclosures, and added a water pond (25 L plastic container, 60 × 39 × 16 cm). We have previously shown that these conditions allowed the discrimination of dispersal events in butterflies (Trochet *et al.*, 2013; Legrand *et al.*, 2015). In this study, we also added a small fenced water pond (similar as the other water pond) in all departure enclosures.

##### Treatments

Twenty damselflies (approx. 40% females) were released in each departure enclosure which represent a density within the range of naturally observed populations (0.2 individual per m^2^, Hardersen 2008). After releasing damselflies and after measuring their activity, we applied the food resources and the predator cues treatments. We used a 2 × 2 factorial design, crossing resource availability (RA) and predation risk (PRED) with two levels each, resulting into 2 departure enclosures of each combination of treatments. The RA and PRED treatment were applied after measuring activity levels and kept for the acclimation phase. During the acclimation phase, which lasted 24 hours, corridors were kept closed to prevent any movements between departure and arrival enclosures. After 24 hours, corridors were opened and we monitored dispersal movements as described below.

##### Resource availability

RA includes two treatments: a low and high resources treatment. In the low food treatment, we kept enclosures with their natural insect community. In the high food treatment, we added approx. 100 fruit flies (mix of *Drosophila* species with body sizes comprised between 2 and 6 mm) per enclosures and a fruit mixture with approx. 200 pupae emerging gradually throughout the experiment. Small Dipterans are the main prey for adult damselflies (Sukhacheva, 1996) and fruit flies were already used as experimental food resources (Debecker *et al.*, 2016). We estimated the relative amount of flying insects after our experiment using a swiping net. The addition of *Drosophila* in the high resources treatment resulted in an increased amount of 3 – 6 mm flying insects (*F*_1,6_ = 5.33, *P* = 0.06, *R*^2^ = 0.47), but not of flying insects smaller than 3 mm (*F*_1,6_ = 0.02, *P* = 0.90). As we did not find any fruit flies in the samples, the effect on 3 – 6 mm flying insects is likely an indirect effect of damselflies eating preferentially fruit flies rather than other flying insects.

##### Predator cues

PRED includes a predation risk treatment, where two frogs were added in a fenced pond, and a no predation risk treatment where the fenced pond was left empty. The addition of frogs produced both a visual, auditive and olfactory cues with no actual predation. Damselflies are known to select oviposition sites depending on the presence of predators (Blaustein, 1999) and our predator treatment mimics this. Enclosures naturally contain several spiders, which are potential predators of damselfly. However, we removed most of them before the experiment and we also estimated their relative abundance after the experiment. We checked that adding this relative abundance in our statistical analyses did not change our results.

##### Dispersal and data collection

After the opening of corridors, we monitored dispersal daily for 5 days. To do so, two people entered and walked quietly for 5 min in each cages. When a damselfly was located, the observer recorded the individual number. This procedure is commonly used on butterfly (Legrand *et al.*, 2012; Trochet *et al.*, 2013; Legrand *et al.*, 2015) and we did not observe any individuals leaving the cage during dispersal monitoring. On the fifth day, we captured all survivors in departure and arrival cages. We attributed a disperser status to individuals which moved at least once between cages and a resident status to individuals which never left the departure cage. We retained individuals which were alive after three days and excluded individuals which were never observed as their dispersal status is unknown.

#### Marsh cricket — *Pteronemobius heydenii*

**Authors** Delphine Legrand, Alexandre Vong and Julien Cote

**License** 2012-10 DREAL (common lizard); 09-2016-02 (common toad)

##### Study organism and predator

The Marsh cricket (*Pteronemobius heydenii*, Fischer 1853) is a small, omnivorous cricket with a brown-black body (males: 5–7 mm, females 6–8 mm) found in humid habitats throughout Eurasia (Iorgu & Iorgu, 2008; Hao-Yu *et al.*, 2016). We used 311 crickets captured in the Metatron (Legrand *et al.*, 2012) and maintained in small tanks (22 L, 39 × 28 × 28 cm). Tanks contained 10 cm of soil litter, 3 egg boxes used as refuges, two small dishes for water and a regular addition of fresh vegetables (apple, grass, potatoes and carrots) after checking that the species effectively fed on them. As we caught crickets in semi-natural enclosures, they have never been exposed to predator cues (see below).

The common lizard (*Zootoca vivipara*) and the common toad (*Bufo bufo*) were used as our model predator. Both species are generalist feeders preying upon various arthropods species including crickets (Avery, 1966; Linzey *et al.*, 1998; Crnobrnja-Isailović *et al.*, 2012). Lizards were raised in cattle tanks as described in the common lizard protocol and toads were captured in the Metatron.

We estimated that crickets had smaller home ranges than lizards and toads (< 10 m^2^ for crickets, roughly estimated from a net squared displacement of a similar sized cricket species, Brouwers & Newton 2010; < 1200 m^2^ for common lizards, (Clobert *et al.*, 1994); 8000–28 000 m^2^ for common toads, Parker & Gittins 1979). Mobility, estimated as sprint speed, was similar for the 3 species (lizards: 30–60 cm s^−1^, Van Damme *et al.* 1991; Sorci *et al.* 1995; toad: approx. 30 cm s^−1^, Beck & Congdon 2000; crickets: 25 cm s^−1^, we estimated the escape speed of crickets using the escape success–predator velocity relationship described in Dangles *et al.* 2006).

##### Experimental setup

We used 8 dispersal systems placed in a greenhouse with controlled temperature (16–25 C). Each system is made of two 130 L plastic containers (78 × 56 × 43 cm) connected by a circuitous plastic pipe (diameter: 10 cm, total length: 4.4 m) on the upper section of the container. The departure container was filled with soil litter providing access to the corridor while we only added a thin layer of soil in corridors and arrival containers. To go from the departure to the arrival containers, crickets had to enter this narrow corridor and fall into the arrival container.

##### Treatments

We created 32 populations made of 8–11 crickets (approx. 9.71 *±* 0.11 SE). We used a 2 × 2 factorial design, crossing resource availability (RA) and predation risk (PRED) with two levels each, resulting into 8 replicates of each combination of treatments. We ran the 8 replicates in 4 blocks, from September 26th to October 20th 2016. The RA and PRED treatments were applied before releasing crickets for a 24 hours acclimation phase with connections between containers closed. After 24 hours, connections were opened and we monitored dispersal movements as described below.

##### Resource availability

RA includes two treatments: a low and high resources treatment. The resources are manipulated through the addition of fresh vegetables (half of a potato, half of a carrot, half of an apple and a handle of grass) in the high resources treatment and only a very small piece of vegetable and 2 pieces of grass in the low food resources. It allows creating a large difference in food availability between high and low food enclosures.

##### Predator cues

Lizards were housed in individual terraria containing 3 cm of soil, a shelter (a piece of eggs carton), a water dish, and a piece of absorbent paper to collect odours serving as predator cues. In one corner of the terrarium, ultraviolet and incandescent lamps provided light and heat for thermoregulation from 9:00 to 12:00 and from 14:00 to 17:00. Lizards were fed daily with 1 cricket (*Acheta domestica*). We used a mix of several absorbent papers belonging to several lizard terraria. Four toads were maintained all together in a large plastic tank (130 L, 78 × 56 × 43 cm) with rocks, water, soil and absorbent paper. Tanks were watered daily and abundantly to maintain enough humidity. Before releasing crickets, dispersal systems with predator cues treatments received a piece of lizard paper mixes and a piece of toad paper, whereas control treatments received a piece of odour free paper collected from vacant containers maintained under the same conditions than inhabited terraria. When removing a piece of paper from predator or predator-free container, we added a new piece of paper for future experimental blocks. The protocol to collect lizard olfactory cues is a standard protocol already used several times to elicit social reactions in lizards (Cote & Clobert, 2007; Teyssier *et al.*, 2014).

##### Dispersal and data collection

After the opening of corridors, we monitored dispersal daily for 5 days. To do so, we captured crickets in the arrival containers without disturbing the departure containers. At the end of the dispersal assay, we captured all crickets in departure and arrival containers. We recaptured 266 crickets over the 311 released, other crickets might have died during the dispersal assays. We only used recaptured crickets in our analysis to prevent confounding dispersal with survival.

#### *Tetrahymena* sp

**Authors** Staffan Jacob, Estelle Laurent and Nicolas Schtickzelle

##### Study organism and predator

We used two species of *Tetrahymena* ciliates (*T. thermophila* and *T. elliotti*), that is, freshwater approx. 50 µm long protists that actively swim and disperse using ciliae beating movements, foraging on bacteria in nature (N-America). They were cultured in the lab under standardized conditions: 23 °C temperature, in a homogeneous nutrient broth (PPYE medium: 2% Proteose peptone and 0.2% yeast extract diluted in ultrapure water) and under strict axenic conditions (sterilized material and culture medium; all manipulations under a flow hood). *Tetrahymena thermophila* has been used as a model species to study different aspects of dispersal: e.g. intraspecific variation (Pennekamp *et al.*, 2014), dispersal syndromes and link with cooperation strategies (Fjerdingstad *et al.*, 2007; Schtickzelle *et al.*, 2009; Chaine *et al.*, 2010; Jacob *et al.*, 2016b), phenotypic plasticity and density dependence (Pennekamp *et al.*, 2014; Jacob *et al.*, 2016b), non-random and informed dispersal (Jacob *et al.*, 2015, 2016b). The predator used was *Blepharisma* sp., a mobile predator previously shown to be an appropriate predator for *Tetrahymena* ciliates (Fronhofer et al. in prep).

##### Experimental setup

The experiments were conducted in standard two-patch systems consisting of two habitat patches (1.5 mL standard microtubes), connected by a corridor (4 mL internal diameter silicon tube, 2.5 cm) and filled with growth media (Fjerdingstad *et al.*, 2007; Schtickzelle *et al.*, 2009; Chaine *et al.*, 2010; Pennekamp *et al.*, 2014; Jacob *et al.*, 2016b,a). Cells were placed in the origin patch and the corridor was opened to allow dispersal towards the target tube. At the end of the dispersal time, the corridor was clamped to separate residents (cells remaining in the origin patch) from dispersers (cells that moved to the target patch).

For each *Tetrahymena* species, we used four isolated genotypes (*T. thermophila* strains: 18282-1, 18282-4, 18296-4, 19876-1; *T. elliotti* strains: 18470-2, 18765-1, 19484-1, 18662-4; Tetrahymena Stock Center, Cornell University, New York) that we mixed at equal density to obtain genetically variable populations (final cell density: 40000 cells mL^−1^). Genotypes were mixed just before the beginning of the experiment, and 200 µL of each mix were inoculated in the start patch of each two-patch system. After one-hour acclimation, corridors were opened to allow cells to disperse for four hours.

##### Treatments

Predator cues (yes/no PRED) and resource availability (standard/low RA) were manipulated in the start patches, with 5 replicates per species and treatment.

##### Resource availability

The low RA treatment consisted in filling the start patch with either standard resource concentration or 10% of standard concentration. The target patch was always filled with standard resource concentration. Furthermore, corridors were filled with water to create an unsuitable matrix containing no resource.

##### Predator cues

The predator cues were obtained by filtering *Blepharisma* sp. culture at carrying capacity (i.e., 2 weeks old culture) using 0.2 µm filters. 100 µL of filtered predator culture, thus containing only chemicals from the predator culture, were then added in the start patch of the two-patch systems for the predator cues treatment. Since predators (*Blepharisma* sp.) were cultured on protozoan pellet (0.46 g L^−1^; Carolina Biological Supply) with bacteria (*Serratia fonticola*, *Bacillus subtilis* and *Brevibacillus brevis*), we performed preliminary analyses using a filtrate from a bacteria culture on protozoan pellet instead of predator cues, and found no effect on dispersal rate compared to ‘no cues’.

##### Data collection

We used a standardized procedure to measure cell density and morphology based on automatic analysis of digital images (Pennekamp & Schtickzelle, 2013). For each culture sample (i.e. a specific tube of a specific 2 patch system), we pipetted five 10 µL samples into a multichambered counting slide (Kima precision cell 301890), and took one digital picture of each chamber under a dark-field microscope (Fjerdingstad *et al.*, 2007; Schtickzelle *et al.*, 2009; Chaine *et al.*, 2010; Pennekamp *et al.*, 2014; Jacob *et al.*, 2015, 2016b,a). Digital pictures were analysed using ImageJ software (version 1.47, National Institutes of Health, USA, http://imagej.nih.gov/ij; Pennekamp & Schtickzelle (2013)) to obtain the overall number of cells on the picture, from which the abundance in each culture tube was recomputed.

#### Two-spotted spider mite — *Tetranychus urticae*

**Authors** Stefan Masier and Dries Bonte

##### Study organism and predator

The model species used for this experiment is the haplodiploid spider mite *Tetranychus urticae* Koch (Acarina: Tetranychidae), also known as two-spotted spider mite, belonging to the ‘LS-VL’ strain. The original population of *T. urticae* was collected in October 2000 from a botanical garden in Ghent (Belgium) where pesticides had not been used for at least ten years (Van Leeuwen *et al.*, 2008).

Mites are phytophagous, aboveground generalist herbivores feeding on plant cell fluids. They are a common pest in gardens, fields and greenhouses and can feed on a large variety of plants species. The bean is well-known from the literature as one of the species the spider mites can feed on most efficiently, as it does not produce harmful chemicals when attacked (opposed to other plants like peppers or tomatoes). The average body length for an adult female is approx. 0.4 mm (Fasulo & Denmark, 2000).

*Phytoseiulus persimilis* Athias-Henriot is well-known as a voracious predator of *T. urticae* and can have a strong impact on the population dynamics of its prey (Sabelis & Van der Meer, 1986; Grostal & Dicke, 1999). The close ecological relation between the two species is widely described (e.g., see Dicke & Sabelis, 1987), and this predator is also commonly used as a pest control in greenhouses. For these reasons, we were reasonably sure we could expect plastic response in the prey behavior when faced with predator cues (Pallini *et al.*, 1999). Avoidance had been the most studied response, even if other reactions have been recently found in closely related species (Lemos *et al.*, 2010). We bought a vial with approx. 200 individuals from a professional seller and we raised them on bean plants infested by *T. urticae* for 1 generation before performing the experiment.

The two-spotted spider mites were raised on an optimal host, the common bean (*P. vulgaris* L., variety ‘Prelude’). The plants were grown in an herbivore-free and pesticide-free walk-in climate room at 25 °C and under a 16/8 h (L/D) photoperiod. Regularly, two-weeks old bean plants were moved to a second climatically controlled room (25 *±* 1 °C) with the same light regime and humidity where the population of mites was kept in open plastic boxes. The same conditions of light and temperature had been maintained for the whole duration of the experiment, except when explicitly noted.

Before the experimental setup, a set of adult females from the stock population of *T. urticae* was synchronized to avoid any possible age-related effect. In order to do so, 50 females had been randomly collected and placed on a 7 × 7 cm square patch freshly cut from a bean leaf. After 48 hours, the females were removed and the leaf, mounted on a bed of moisturized cotton to keep it fresh, was placed into a heated cabinet to prevent contamination and provide optimal conditions for the development of the juveniles. The eggs were monitored daily until they reached the adult stage (approx. 7 days at 30 °C). We chose to select for the experiment only mated, one-to-two days old females of *T. urticae*, as they are well-known from the literature as the main dispersing stage in the mites life cycle (Kennedy & Smitley, 1985; Li & Margolies, 1993).

##### Experimental setup

To test for dispersal propensity, the two-patches system was composed as following. A 2.5 × 1.5 cm leaf square cut from a two-weeks-old plant of *P. vulgaris* have been mounted on a cotton bed and fixed in place using paper stripes. The cotton and the paper are to be constantly kept moist to avoid dehydration of the leaf and to prevent the mites from escaping. The paper strips also provided the additional value to stop the mites from crawling under the leaf, where they would have been at risk of drowning, and to protect the exposed edges of the leaf from dehydration. The patch of origin was then connected to an identical target patch using a ParafilmⓇ bridge, that constitutes an unsuitable environment for the spider mites. The length of the bridge had been set at 8 cm, as during pilot experiments this distance was shown to set a dispersal rate between 15% and 20%.

##### Treatments

The population on the patch of origin was composed by one-to-two days old adult, fertilized female spider mites, to maximize the number of individuals suitable for dispersal and subsequent analysis. All the individuals used for the experiment belong to the LS-VL strain. On the patch of origin, 50 individuals were placed, for a density of approx. 13 mites cm^−2^. This mimics a mild-to-severe infestation in natural conditions, and can stimulate the dispersion of individuals (Bitume *et al.*, 2013).

The patches or origin presenting the predator cues (yes PRED) were prepared by letting a sample of 5 predatory mites wander freely on the leaf cut for 24 hours right before the beginning of the experiment. This number was chosen to ensure abundant cues of the presence of *P. persimilis* were left on the leaf: it was shown that two-spotted spider mites are able to spot this amount of predators when present on a leaf and modify their behavior accordingly (Kriesch & Dicke, 1997). The predators were well-fed before being moved to the experimental patch, as no prey was present on the leaf to avoid confounding effects due to individuals feeding on the leaf, and thus potentially changing the starting condition of the patch, or leaving information on the previous population composition, eventually including relatedness. All the alive predator mites and eggs were then removed before placing the experimental population, leaving only the webs and the fecal material laid during their stay. Only patches presenting at least three alive predators and/or one egg were chosen for the experiment, to ensure that predators spent a significant amount of time on the leaf. For the removal, a thin paint brush was used; special attention was paid to ensure no contamination between the patches exposed to predators and the control ones by carefully cleaning the hairbrush with water and ethanol after each patch preparation.

We left the mites settle for 24 hours before connecting the bridges, and then left the spider mites free to disperse for the following 48 hours. We ran five different replicates, each one presenting the four different treatments at once. The replicas are independent one from the other as each time new individuals belonging to the synchronized offspring of stock females were used.

##### Resource availability

From previous experiments, we assessed that a 2.5 × 1.5 cm leaf square cut from a fresh bean leaf is an abundant source of food that can sustain approx. 50 mites for at least 48 hours. As such, this kind of setting was used for the standard RA experiments. The leaves from two-week-old bean plants were cut from the plant, shaped accordingly and left on a wet layer of cotton for 48 hours to ensure hydration before the start of the experiment.

During the low RA treatment, the bean leaves underwent the same treatment, but a square of ParafilmⓇ was placed between the cotton bed and the leaf to isolate the leaf itself from the water. Doing so, the leaves withered and lost wet weight, presenting less food available for the animals to eat, as mites are not able to feed on plant tissues with a reduced percentage of water. The cotton had to be kept wet in the predator treatments to prevent individuals from escaping, so water was added to all the preparations to maintain the humidity levels constant through all the replicas and avoid adding an unwanted difference between the two low RA treatments. Pilot choice tests were performed in advance showing that, while mites do not significantly discriminate between fresh and 24-hours-old withered leaves, they show a marked avoidance for leaves that were left to wither for more than 48 hours.

The amount of food consumed by mites can be measured by counting the feeding scars left on the leaves and comparing the scarred area with the patch total area (Kant *et al.*, 2004). This procedure had been largely applied in previous studies and it can be standardized using specific software tools. Before and after the experiment, a picture of every patch was taken and the pairs were digitally compared to evaluate the consumption rate and the per capita amount of consumed resource.

##### Predator cues

Mites are known to be able to detect the presence of a predator mostly using chemical cues. In particular, hormones secreted by the predators seem to be the major cue for the spider mites to avoid leaves infested by predators (Kriesch & Dicke, 1997; Dicke & Grostal, 2001). As the leaves were directly exposed to the predators for 24 hours right before the experiment, the risk of interference due to solvents and/or other chemicals used for the extraction is not present in this design.

##### Data collection

After dispersal time is over, the bridges were removed and the individuals crawling on both patches and the bridge itself were visually counted. A partial count was also performed roughly every 24 hours to ensure some intermediate data as well.

#### Common lizard — *Zootoca vivipara*

**Authors** Laurane Winandy, Félix Pellerin, Lucie di Gesu, Delphine Legrand and Julien Cote

**License** 2012-10 DREAL

##### Study organism and predator

The common lizard (*Zootoca vivipara*; Jacquin 1787) is a small lacertid (adult snout–vent length: males, 40–60 mm; females, 45–75 mm) generally found in humid habitats throughout Eurasia. In this experiment, we used 112 two-year old lizards born in the lab (Ariège, France) and raised in 1100 L cattle tanks (diameter: 1.70 m). Home tanks contained 20 cm of soil litter, dense vegetation, ten 50 mL Falcon tube in the litter and 3 half flower pots used as refuges, two small dishes for water and a regular addition of crickets. These conditions were highly suitable for lizards, as shown in previous experiments (Teyssier *et al.*, 2014; Bestion *et al.*, 2014, 2015). As lizards spent their entire life in cattle tanks, they have never been exposed to real predators. Individuals were all marked at birth (see Bestion *et al.*, 2014), sexed, and before the experiment, we measured several phenotypic traits (see below). After measurements, individuals were transferred to our experiment dispersal systems (see below).

The Green whip snake (*Hierophis viridiflavus*) was used as our model predator. Adult green whip snakes are generalist feeders, preying upon small mammals, reptiles and birds, but neonates are highly specialized on lizards (Lelièvre *et al.*, 2010). Green whip snakes occur in sympatry with common lizards in their southern distribution. We used two snakes successively to prevent maintaining a single individual over a long period of time. We maintained snakes in our laboratory in a room separated from the lizards’ room.

We estimated that lizards have a smaller home range than the green whip snake (< 1200 m^2^ for common lizards, Clobert *et al.* 1994, *>* 10 000 m^2^ for snakes, estimated from Lelièvre *et al.* 2010), while the mobility, estimated as sprint speed, was higher for snakes than for lizards (lizards: 30–90 cm s^−1^, Van Damme *et al.* 1991; Sorci *et al.* 1995; snakes: 15–90 cm s^−1^, Lelièvre *et al.* 2010). On top of dispersal, tail autotomy is another defense against predators (Bateman & Fleming, 2009).

##### Experimental setup

We used 8 dispersal systems made of connected 1100 L cattle tanks similar to home tanks. These systems are made of 4 cattle tanks juxtaposed and connected by a plastic pipe (diameter: 20 cm). In total, systems are 7 m long. The first and last cattle tanks contained the same environments as home tanks. To go from the first to the last tank, lizards had to cross two other tanks containing 20 cm of soil covered with a fake road made of tarmac roofing free of shelters. These two intermediate tanks simulated a 3.4 m wide road making movements between naturalized tanks. In the arrival tank, we removed soil underneath the connecting pipe to prevent dispersing lizards moving back to their initial tank.

##### Treatments

We created 16 populations made of 7 lizards (3 females and 4 males for 10 populations and 2 females and 4 males for 6 populations). The 7 lizards were released in departure tanks which represent a density within the range of naturally observed populations (Bestion *et al.*, 2015). We used a 2 × 2 factorial design, crossing resource availability (RA) and predation risk (PRED) with two levels each, resulting into 4 departure enclosures of each combination of treatments. We ran the four replicates in 2 blocks, one on July 16th and one on August 2nd. The RA and PRED treatments were applied before releasing lizards in departure tanks. After a 24 hours acclimation phase with connections between tanks closed, connections were opened and we monitored dispersal movements as described below.

##### Resource availability

RA includes two treatments a low and high resources treatment. The resources are manipulated through the addition of 200 crickets in the high resources treatment. Cattle tanks had some natural resources for lizards to survive in the low resources treatment. We estimated the relative abundance of spider and orthoptera, the main prey for common lizards (Avery, 1966). Our treatments created a large difference in food availability between high and low food enclosures (*F*_1,14_ = 27.52, *p* = 0.0001).

##### Predator cues

The snake was kept in a separate room in a wood box (50 × 40 × 10 cm) featuring a clean water bowl, a hiding spot and a light bulb for basking (40 W; set on a 12 L: 12 D cycle). In order to collect snake odors, we placed 40 small calcite tiles (3 × 3 × 0.6 cm) in the snake cage. The tiles were left 3 days before being transferred into the dispersal systems. Upon collection, tiles were gently rubbed against the snake belly and sides in order to saturate them with snake odour. Forty identical tiles, kept in a separate room, were used as control for the predator free treatment. Before releasing lizards, dispersal systems with predator cues treatments received 10 tiles collected from the snake terrarium, whereas control treatments received control, odour-free tiles. After each assay, all slabs were cleaned with 70% ethanol, rinsed and put back with snakes or in a control box. This procedure has been repeatedly shown to efficiently elicit antipredator responses (Teyssier *et al.*, 2014; Bestion *et al.*, 2014, 2015).

##### Dispersal and data collection

After the opening of corridors, we monitored dispersal daily for 9 days. To do so, we captured lizards in the arrival tanks without disturbing the departure tanks. At the end of the dispersal assay, we captured all lizards in departure and arrival tanks. Ten lizards were found in the intermediate tanks and, as we were unsure of their dispersal status, we excluded them from the analysis

**Figure S1:**
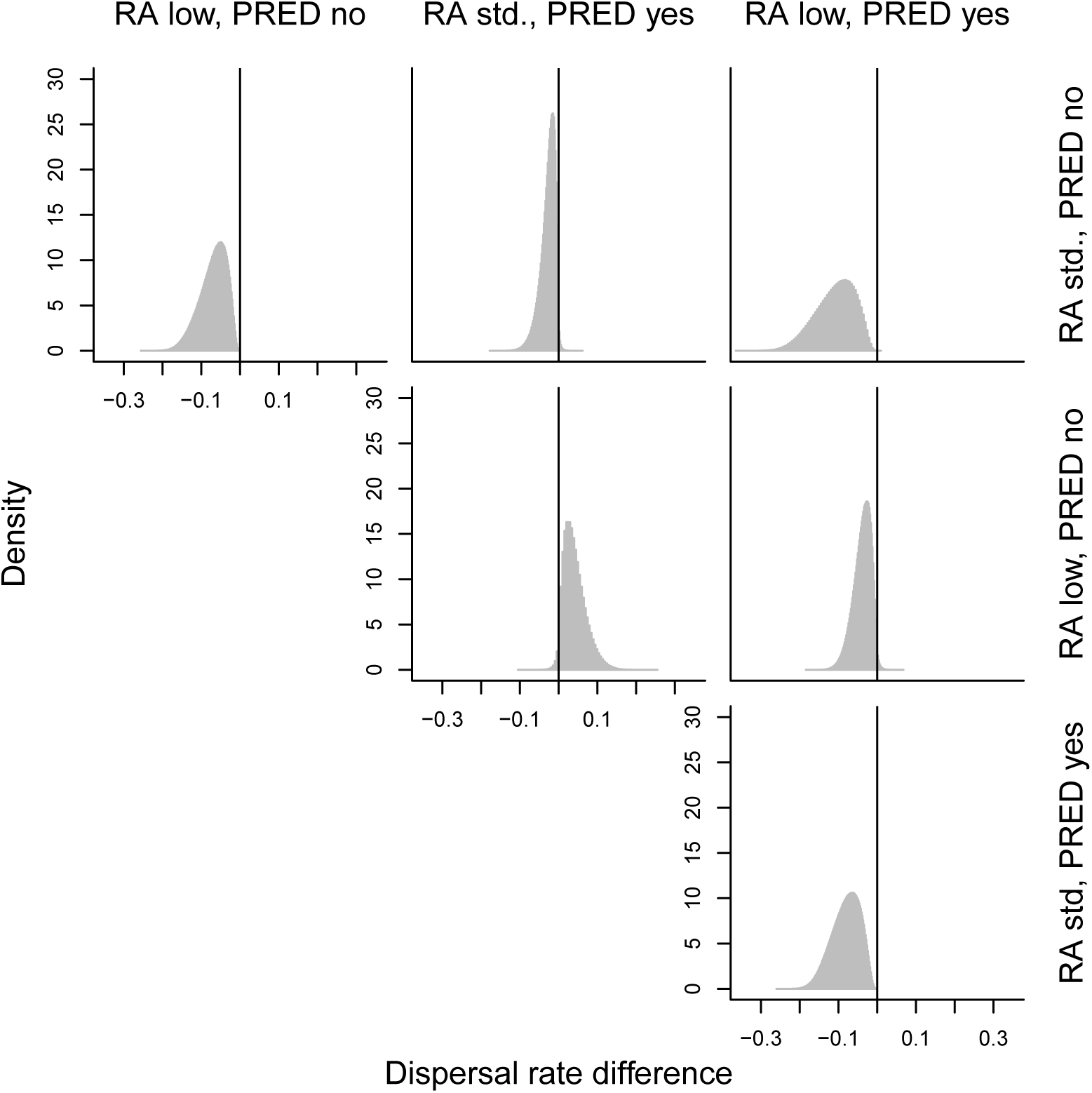
Pairwise differences between posterior predictive distributions of dispersal rates (back-transformed) in the four treatments for the most parsimonious, that is, additive model.

**Figure S2:**
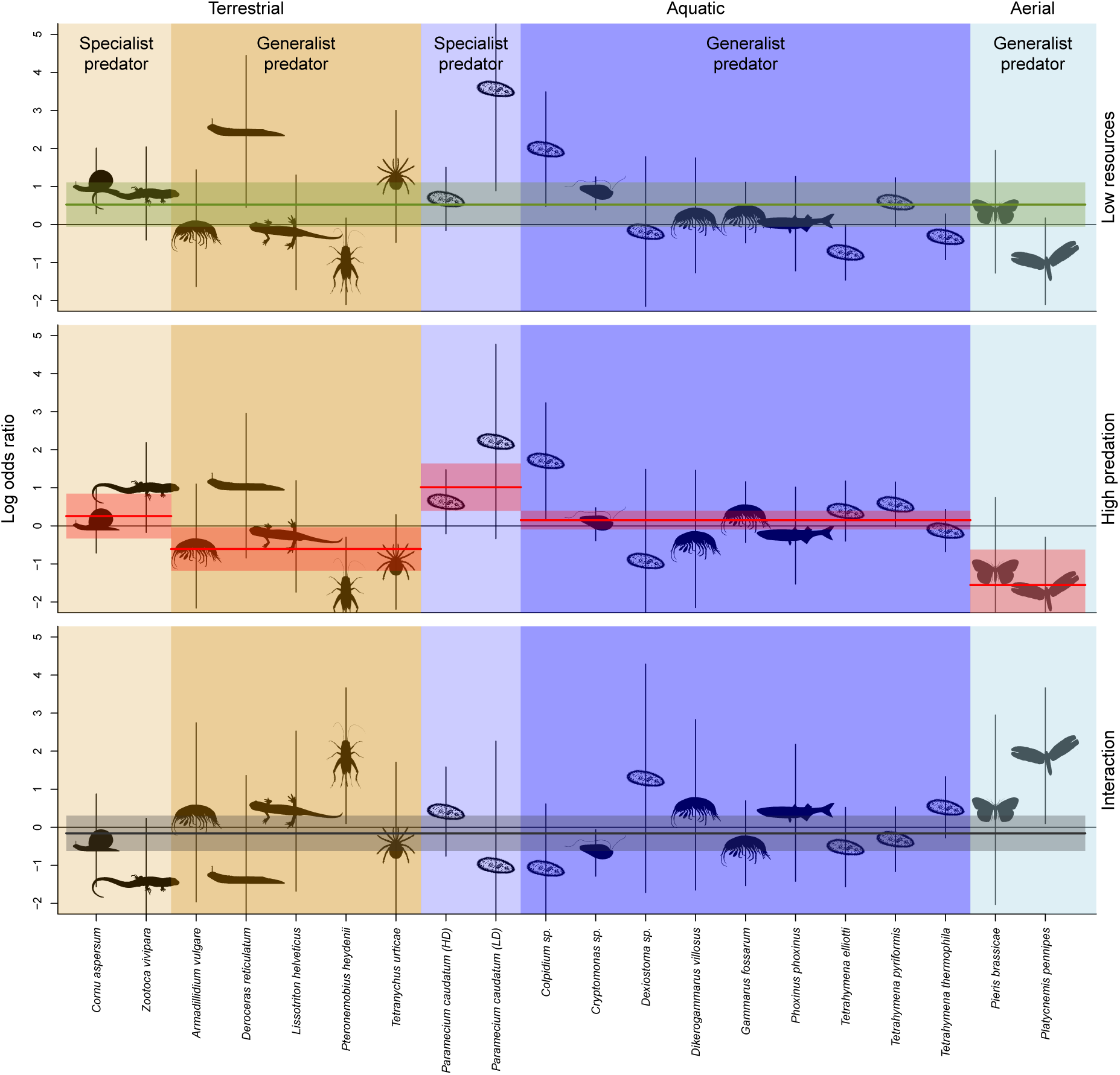
Effect of resource limitation, predation risk and the interaction between these two factors on dispersal for all study species individually. We show log odds ratios (logOR; black animal symbols) and confidence intervals (vertical black lines) of the resource limitation (top), predation risk (center) and interaction (bottom) effects per species sorted according to dispersal mode. Model predictions of the model retained after model selection (mean and CI) are superimposed. See Tab. S6 for model selection results.

**Figure S3:**
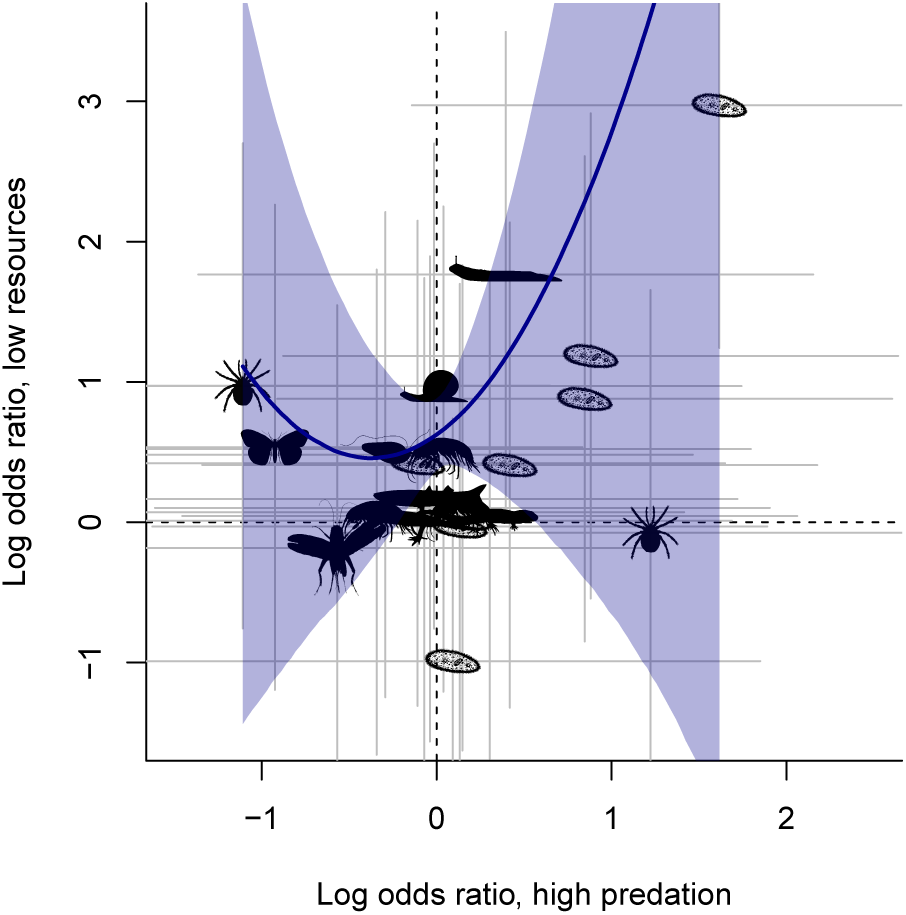
Correlation of responses to resource limitation and predation risk. Animal symbols and corresponding grey lines show log odds ratios and corresponding confidence intervals per species. The solid blue line and the shaded area visualize the quadratic relationship between the log odds ratios suggested by model selection (see main text).

**Figure S4:**
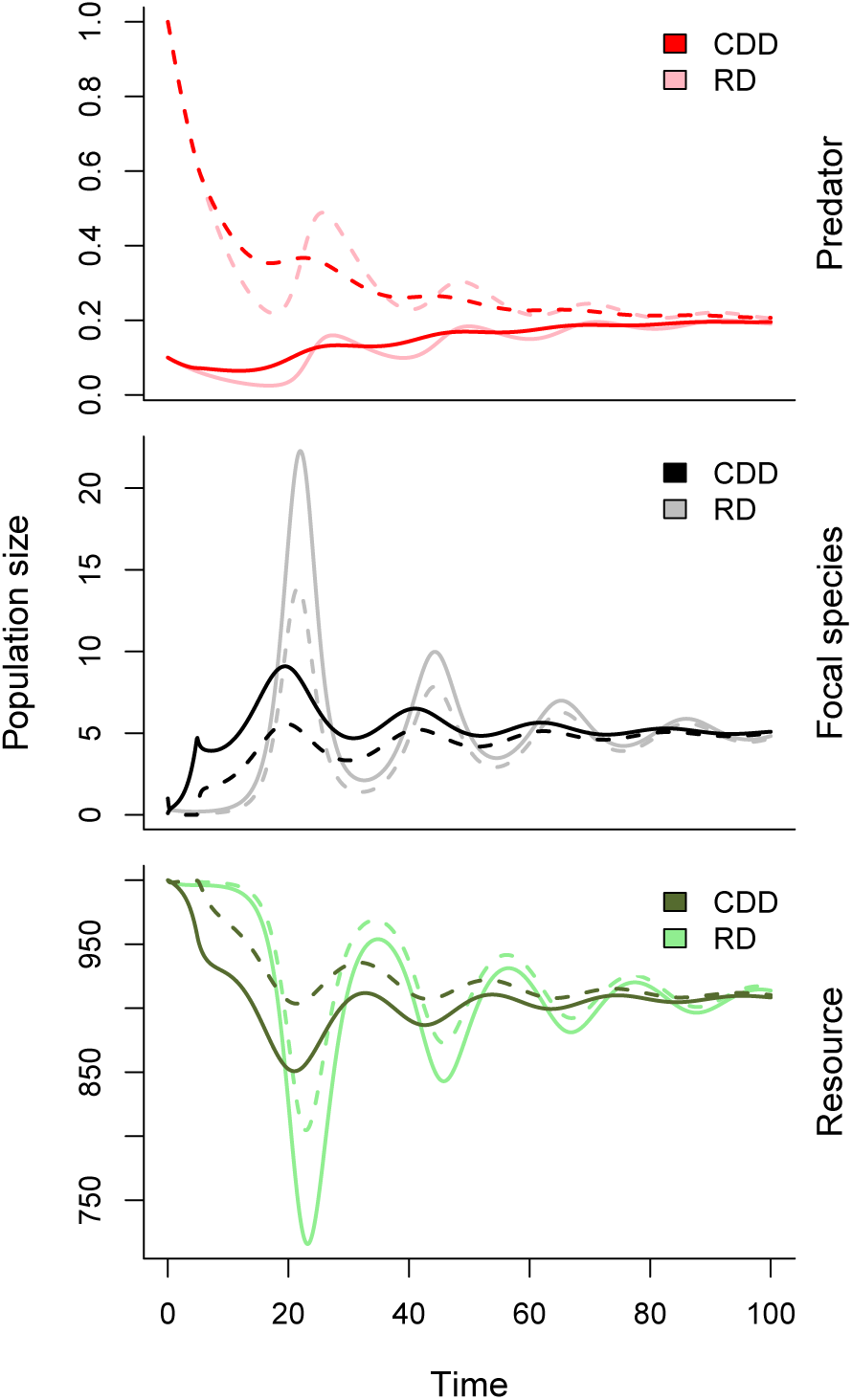
Consequences of context-dependent dispersal for population and metacommunity dynamics. We show the dynamics of all three tropic levels (resources, *R*; focal species, *N*; top predator *P*) in both patches (patch 1: solid lines, patch 2: dashed lines). While the RD and CDD scenarios are characterized by the same model parameters, we compare the specific scenarios in which the CDD parameters minimize the focal species population dynamics CV (as in the main text; *T_R_* = 956.94 and *T*_*P*_ = 0.12) with the RD scenario that exhibits the same dispersal rate at population dynamic equilibrium (i.e, *m*_*N*_ = 1). The results are not qualitatively changed, except for the covariance in predator dynamics: the CV of the focal species (resource, predator) population dynamics is reduced by 49% (48%, 8%) in the CDD scenario compared to RD while the covariance between dynamics in patches 1 and 2 is reduced by 88% (80%, increased by 48%). Parameter values: ω = 0.5, *R*_0_ = 1000, *e*_*N*_ = 0.1, *a*_*N*_ = 0.01, *d*_*N*_ = 0.1, *e*_*P*_ = 0.005, *a*_*P*_ = 4,*d*_*P*_ = 0.1.

**Table S1:**
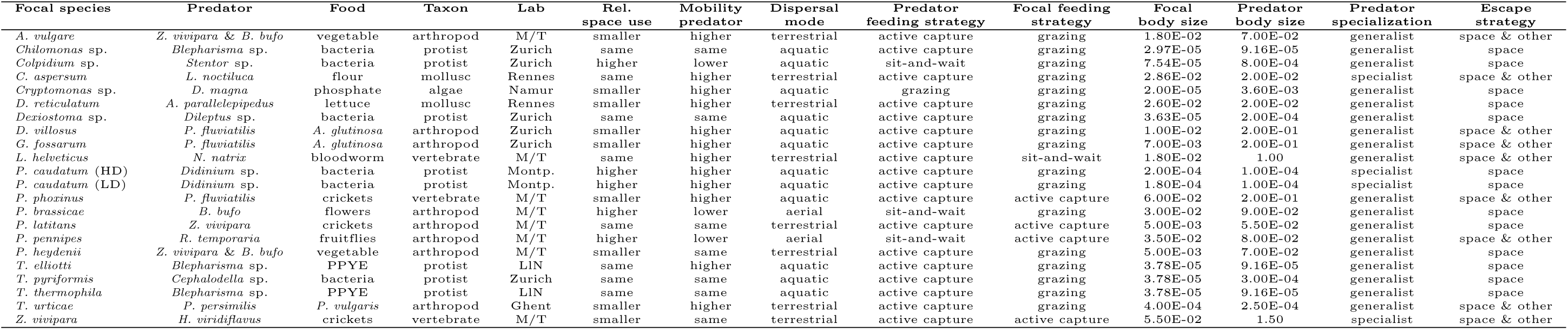
Table summarizing the species specific information. Laboratory abbreviations: Montpellier (Montp.), Moulis/ Toulouse (M/T), Louvain-la-Neuve (LlN). Body size is indicated in metres. Feeding strategies were classified according to Dell *et al*. (2014). If not indicated otherwise in the respective methods section, data contained within this table was measured during the experiment. For qualitative information, such as relative space use or mobility, we were conservative, ignoring differences roughly within one order of magnitude.

**Table S2:** Model selection results for the overall effect of resource limitation (RA) and predation risk (PRED). For a visualization see Fig. 1.

| **Model** | **$\Delta\text{AICc}$** | **$W_{\text{AICc}}$** |
| --- | --- | --- |
| RA + PRED | 0 | 0.62 |
| RA * PRED | 1.79 | 0.25 |
| RA | 3.13 | 0.13 |
| PRED | 24.19 | 0 |
| 1 | 27.15 | 0 |

**Table S3:** Model selection results for the effect of resource limitation (RA) — additive model. We only show the top five models. For a visualization see Fig. 2.

| Model | $\Delta\text{AICc}$ | $W_{\text{AICc}}$ |
| --- | --- | --- |
| 1 | 0 | 0.75 |
| log(focal body size) | 2.94 | 0.17 |
| focal feeding strategy | 6.34 | 0.03 |
| dispersal mode | 6.51 | 0.03 |
| log(focal body size) + focal feeding strategy | 10.29 | 0 |

**Table S4:** Model selection results for the effect of predation risk (PRED) — additive model. We only show the top ten models. For a visualization see Fig. 2.

| Model | $\Delta\text{AICc}$ | $W_{\text{AICc}}$ |
| --- | --- | --- |
| dispersal mode + rel. space use | 0 | 0.20 |
| 1 | 0.27 | 0.17 |
| predator specialization | 0.55 | 0.15 |
| dispersal mode + rel. space use + log(focal body size) | 2.46 | 0.06 |
| predator specialization + escape strategy | 2.70 | 0.05 |
| escape strategy | 3.10 | 0.04 |
| log(focal body size) | 3.29 | 0.04 |
| dispersal mode + predator specialization | 3.48 | 0.03 |
| predator specialization + log(focal body size) | 3.75 | 0.03 |
| dispersal mode + predator specialization + predator feeding strategy | 4.15 | 0.02 |

**Table S5:**
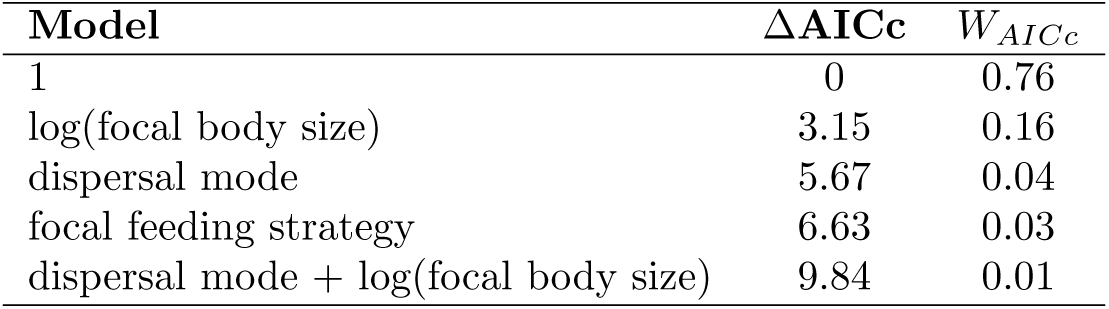
Model selection results for the effect of resource limitation (RA) — interaction model. We only show the top five models. For a visualization see Fig. S2.

| Model | $\Delta AICc$ | $W_{AICc}$ |
| --- | --- | --- |
| 1 | 0 | 0.76 |
| log(focal body size) | 3.15 | 0.16 |
| dispersal mode | 5.67 | 0.04 |
| focal feeding strategy | 6.63 | 0.03 |
| dispersal mode + log(focal body size) | 9.84 | 0.01 |

**Table S6:** Model selection results for the effect of predation risk (PRED) — interaction model. We only show the top ten models. For a visualization see Fig. S2.

| Model | $\Delta AICc$ | $W_{AICc}$ |
| --- | --- | --- |
| dispersal mode + predator specialization | 0 | 0.38 |
| 1 | 2.46 | 0.11 |
| predator specialization | 2.58 | 0.11 |
| dispersal mode | 2.97 | 0.09 |
| predator specialization + log(focal body size) | 4.78 | 0.04 |
| dispersal mode + predator specialization + log(focal body size) | 4.83 | 0.03 |
| dispersal mode + predator specialization + escape strategy | 4.84 | 0.03 |
| escape strategy | 5.38 | 0.03 |
| log(focal body size) | 5.61 | 0.02 |
| predator specialization + escape strategy | 6.16 | 0.02 |

**Table S7:** Model selection results for the effect of predation risk (interaction) — interaction model: interaction model. We only show the top ten models. For a visualization see Fig. S2.

| Model | $\Delta AICc$ | $W_{AICc}$ |
| --- | --- | --- |
| 1 | 0 | 0.32 |
| log(focal body size) | 2.35 | 0.10 |
| predator specialization | 2.77 | 0.08 |
| dispersal mode | 2.78 | 0.08 |
| escape strategy | 3.17 | 0.06 |
| predator feeding strategy | 3.17 | 0.06 |
| escape strategy + log(focal body size) | 4.46 | 0.03 |
| rel. space use | 4.55 | 0.03 |
| predator specialization + log(focal body size) | 5.09 | 0.02 |
| predator mobility | 5.27 | 0.02 |

**Table S8:** Sensitivity analysis of the consequences of CDD in the two-patch food-chain model (Eqs. 1a–c). We here report the relative change in coefficient of variation (CV) of the temporal dynamics if dispersal is assumed to be context-dependent instead of random, that is, negative values indicate a reduction in CV in the CDD dynamics compared to RD. We always compare the RD-CDD scenario pair that both minimize the respective CV of the focal species time series. The table shows relative changes for the focal species as well as in brackets for the resource (first value) and for the predator (second value).

| Model parameter | - 50 % of standard value | + 50 % of standard value |
| --- | --- | --- |
| $\omega$ | - 43 % (- 26 %; + 4 %) | - 47 % (- 42 %; + 1 %) |
| $e_N$ | — | - 37 % (- 29 %; - 2 %) |
| $a_N$ | - 30 % (- 26 %; - 4 %) | - 39 % (- 28 %; - 1 %) |
| $e_P$ | - 45 % (- 29 %; + 7 %) | - 43 % (- 39 %; - 6 %) |
| $a_P$ | - 37 % (- 28 %; - 5 %) | - 43 % (- 36 %; - 3 %) |

**Table S9:** Sensitivity analysis of the consequences of CDD in the two-patch food-chain model (Eqs. 1a–c). We here report the relative change in covariance of the temporal dynamics in both patches if dispersal is assumed to be context-dependent instead of random, that is, negative values indicate a reduction in covariance in the CDD dynamics compared to RD. We always compare the RD-CDD scenario pair that both minimize the respective CV of the focal species time series. The table shows relative changes for the focal species as well as in brackets for the resource (first value) and for the predator (second value).

| Model parameter | - 50 % of standard value | + 50 % of standard value |
| --- | --- | --- |
| $\omega$ | - 73 % (- 44 %; - 23 %) | - 84 % (- 78 %; - 6 %) |
| $e_N$ | — | - 82 % (- 70 %; + 7 %) |
| $a_N$ | + 527 % (+ 173 %; - 31 %) | - 94 % (- 73 %; + 3 %) |
| $e_P$ | - 61 % (- 42 %; - 20 %) | - 100 % (- 91 %; - 8 %) |
| $a_P$ | - 74 % (- 58 %; + 26 %) | - 94 % (- 81 %; - 21 %) |

